# Membrane-mediated dimerization potentiates PIP5K lipid kinase activity

**DOI:** 10.1101/2021.09.21.461304

**Authors:** Scott D. Hansen, Albert A. Lee, Jay T. Groves

## Abstract

The phosphatidylinositol 4-phosphate 5-kinase (PIP5K) family of lipid modifying enzymes generate the majority of phosphatidylinositol 4,5-bisphosphate (PI(4,5)P_2_) lipids found at the plasma membrane in eukaryotic cells. PI(4,5)P_2_ lipids serve a critical role in regulating receptor activation, ion channel gating, endocytosis, and actin nucleation. Here we describe how PIP5K activity is regulated by cooperative binding to PI(4,5)P_2_ lipids and membrane-mediated dimerization of the kinase domain. In contrast to constitutively dimeric phosphatidylinositol 5-phosphate 4-kinase (PIP4K, type II PIPK), solution PIP5K exists in a weak monomer-dimer equilibrium. PIP5K monomers can associate with PI(4,5)P_2_ containing membranes and dimerize in a protein density dependent manner. Although dispensable for PI(4,5)P_2_ binding and lipid kinase activity, dimerization enhances the catalytic efficiency of PIP5K through a mechanism consistent with allosteric regulation. Additionally, dimerization amplifies stochastic variation in the kinase reaction velocity and strengthens effects such as the recently described stochastic geometry sensing. Overall, the mechanism of PIP5K membrane binding creates a broad dynamic range of lipid kinase activities that are coupled to the density of PI(4,5)P_2_ and membrane bound kinase.

## INTRODUCTION

Phosphatidylinositol phosphate (PIP) lipids are an important class of second messengers that regulate the localization and activity of proteins on every intracellular membrane in eukaryotic cells (Di Paolo and De Camilli 2006; Balla 2013). Synthesis of PIP lipids is regulated by different classes of lipid kinases and phosphatases that drive the interconversion between PIP lipid species through the phosphorylation and dephosphorylation of inositol head groups. Of particular importance to a vast array of signaling pathways are phosphatidylinositol 4,5-bisphosphate [PI(4,5)P_2_] lipids, which comprise a minor phospholipid component of the total cellular membrane composition (i.e. ~2 %) (Wenk et al. 2003; Nasuhoglu et al. 2002). PI(4,5)P_2_ serves many important functions in biological processes, including receptor activation, ion channel function (S. B. Hansen 2015), endocytosis (Zoncu et al. 2007; Jost et al. 1998), and actin network assembly at the plasma membrane (Janmey, Bucki, and Radhakrishnan 2018). Understanding the mechanisms that control PI(4,5)P_2_ lipid synthesis is critical for deciphering how cells regulate receptor signaling and PIP lipid homeostasis at the plasma membrane.

Two classes of lipid kinases catalyze the production of PI(4,5)P_2_ lipids: phosphatidylinositol 4-phosphate 5-kinase (PIP5K, type I PIPK) and phosphatidylinositol 5-phosphate 4-kinase (PIP4K, type II PIPK) (Burke 2018). These two families of lipid kinases differ in substrate specificity; PIP5K phosphorylates phosphatidylinositol 4-phosphate – PI(4)P, while PIP4K phosphorylates phosphatidylinositol 5-phosphate – PI(5)P (Loijens and Anderson 1996; Rameh et al. 1997; Muftuoglu et al. 2016). Due to the higher abundance of PI(4)P in the plasma membrane, relative to PI(5)P, the majority of PI(4,5)P_2_ lipids are generated from reactions catalyzed by PIP5K (Doughman, Firestone, and Anderson 2003; Balla 2013). Functionally conserved across eukaryotes, yeast express a single PIP5K enzyme – Mss4 (Yoshida et al. 1994; Homma et al. 1998; Desrivières et al. 1998), while three paralogs – PIP5KA, PIP5KB, PIP5KC – regulate the production of PI(4,5)P_2_ lipids in mammalian cells.

Several mechanisms regulate membrane docking of PIP5K, including substrate recognition (Muftuoglu et al. 2016; Kunz et al. 2000), electrostatic interactions with anionic lipids at the cell plasma membrane (Fairn et al. 2009), and membrane insertion of an amphipathic helix (Liu et al. 2016). Single molecule characterization of human PIP5KB has revealed a role for PI(4,5)P_2_ lipids in controlling cooperative membrane association and positive feedback during PI(4)P lipid phosphorylation reaction (S. D. Hansen et al. 2019). Structural and biochemical studies indicate that, like PIP4K (Rao et al. 1998; Burden et al. 1999), PIP5K can homodimerize in solution (Hu et al. 2015). In the case of zebrafish PIP5KA (zPIP5KA), dimerization has been shown to be required for lipid kinase activity (Hu et al. 2015). It remains unclear whether dimerization regulates membrane docking, ATP binding, or catalysis of PIP5K. Overall, the sequence of molecular interactions that control PIP5K membrane localization and activation has not been elucidated.

Using TIRF microscopy to measure the kinetics of PIP lipid phosphorylation on supported lipid bilayers (SLBs) *in vitro*, we previously reported that human PIP5KB catalyzes the phosphorylation of PI(4)P with a positive feedback loop based on association with it’s reaction product, PI(4,5)P_2_ (S. D. Hansen et al. 2019). Based on the crystal structure of zebrafish PIP5KA (Hu et al. 2015; Muftuoglu et al. 2016) and previous biochemical data, we have worked under the assumption that PIP5K functions as an obligate dimer. However, through comparative single molecule characterization of PIP4K and PIP5K membrane binding dynamics *in vitro* we discovered that members of the PIP5K protein family exist in a weak monomer-dimer equilibrium in solution. At low molecular densities, PIP5K can associate with PI(4,5)P_2_ membranes as a monomer and catalyze the phosphorylation of PI(4)P. Under these conditions, the mechanism of PIP5K positive feedback is controlled by cooperative binding to the reaction product, PI(4,5)P_2_. Increasing the surface density of membrane bound PIP5K promotes dimerization, which further increases the dwell time and enhances the catalytic efficiency of the kinase ~20-fold. Consistent with a mechanism of allosteric regulation, dimerization increases PIP5K catalytic efficiency independently of enhancing membrane avidity. We find that the increase in kinase activity afforded by membrane-mediated dimerization—more specifically the strong positive feedback it creates—dramatically enhances the PIP5K’s ability to form bistable PIP compositional patterns in the presence of a 5’-phosphatase on supported lipid bilayers. The membrane-mediated dimerization also amplifies stochastic fluctuations in kinase reaction velocity. In the context of spatial confinement, these magnified fluctuations facilitate mechanisms such as the recently reported stochastic geometry sensing, in which bistability and even the deterministic outcome of a competitive reaction may depend on system size (S. D. Hansen et al. 2019) (in review; A. A. Lee et al., 2021). Together, our results highlight a mechanism by which PI(4,5)P_2_ binding and membrane-mediated dimerization create a broad dynamic range of PIP5K activities that cells can potentially leverage to tune the concentration and spatial distribution of PI(4,5)P_2_ lipids on cellular membranes.

## RESULTS

### PIP4K and PIP5K bind to membranes with distinct oligomerization states

The PIP4K and PIP5K families of lipid kinases reportedly form homodimeric complexes with structurally distinct dimer interfaces (**Figure 1A**; Hu et al. 2015; Rao et al. 1998; Burden et al. 1999). To determine the role dimerization serves in regulating membrane association and activation of PIP5K, we compared the dynamics of PIP4K and PIP5K dimerization in solution and on supported membranes. Consistent with published results (Hu et al. 2015), we found that zebrafish PIP5KA (zPIP5KA), wild-type and dimer mutant, displayed distinct elution profiles by size exclusion chromatography (SEC) (**Figure 1B**). Comparing the elution profiles of zPIP5KA and PIP4KB – both predicted to have dimeric molecular weights of 90 kDa – revealed that zPIP5KA eluted significantly slower compared to PIP4KB (**Figure 1B**). Differences in the elution profile of PIP4KB and zPIP5KA could be the result of distinct subunit orientations or a difference in the oligomerization state (**Figure 1A**). The observed difference led us to test the hypothesis that PIP5K exists in a weak monomer-dimer equilibrium, rather than being a constitutive dimer.

**Figure 1.**
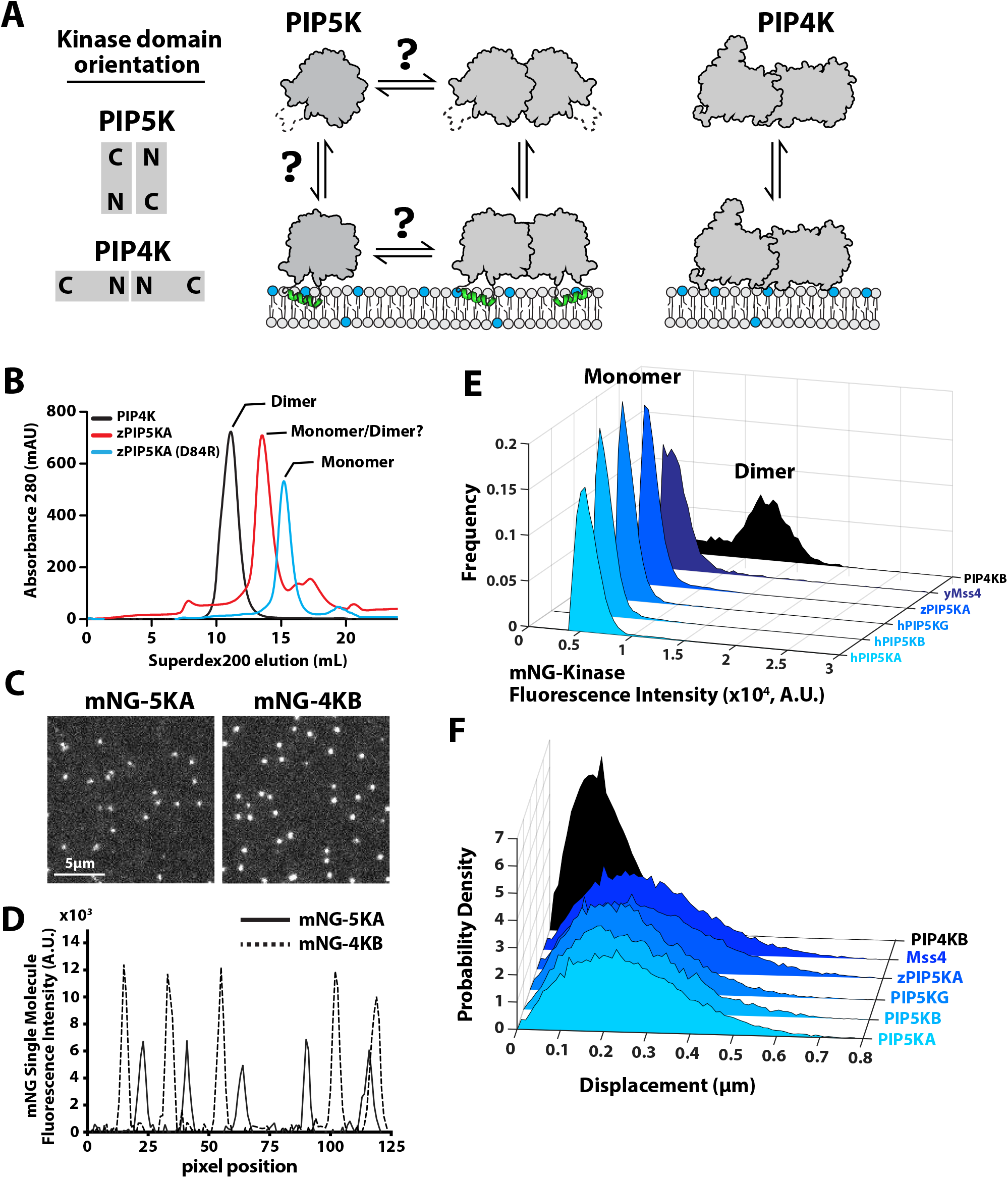
PIP4K and PIP5K bind to PI(4,5)P_2_ membranes with distinct oligomeric states. **(A)** Cartoon showing the orientation PIP4K and PIP5K homodimer subunits relative to the membrane and their proposed membrane bound oligomerization state. (B) Size exclusion chromatography of elution profiles for PIP4KB, zPIP5KA, and zPIP5KA (D51R, dimer interface mutant). (C) Single molecule TIRF microscopy images of IpM mNG-PIP4KB and 5 pM mNG-PIP5KA bound to supported membranes containing 4% PI(4,5)P_2_. **(D)** Intensity line scans of mNG-PIP4KB and mNG-PIP5KA molecules bound to membranes in (C). (E) Frequency distributions of single particle intensities measured for membrane bound mNG-PIP4K and different mNG-PIP5K protein family members. (F) Step size distributions for membrane bound mNG-PIP4K (0.04 μm^2^/sec) and mNG-PIP5KA, PIP5KB, PIP5KG, zPIP5KA, Mss4 (0.14-0.18 μm^2^/sec).

To determine whether PIP4K and PIP5K in solution can be recruited to membranes with distinct oligomerization states, we first established a single molecule cell lysate assay (Lee et al. 2017) to compare the membrane binding properties of single fluorescently labeled PIP4K and PIP5K bound to supported lipid bilayers (SLBs) containing PI(4,5)P_2_ lipids. mNeonGreen (mNG) labeled PIP4K and PIP5K proteins were transiently expressed in human kidney embryo 293 (HEK 293) cells. Cells were lysed by sonication and centrifuged to remove membranes and debris. Using a purified mNG standard, we quantified the concentration of mNG-PIP4K and mNG-PIP5K in clarified cell lysate (**Figure 1–figure supplement 1A**). Samples containing mNG-kinase were diluted in assay buffer to a concentration of 10-100 pM and incubated with supported membranes containing 4% PI(4,5)P_2_ lipids. The resulting surface density of membrane bound mNG-PIP4K and mNG-PIP5K was ~0.03 molecules/μm^2^ (or ~ 100 molecules per field of view).

To quantify differences in the oligomerization states of membrane bound mNG-PIP4K and mNG-PIP5K, we compared the single molecule fluorescence brightness and diffusion coefficients using Total Internal Reflection Fluorescence (TIRF) microscopy (**Figure 1C-1F**). Intensity line scans through mNG foci revealed that the majority of mNG-PIP4KB molecules were two times brighter than membrane bound mNG-PIP5KA (**Figure 1D**). Because mNG-PIP4K is a constitutive dimer, the molecular brightness distribution of this lipid kinase established the upper limit for the percentage of detectable dimers in our assay. This was based on the fraction of mNG molecules that formed fully mature chromophores during expression in HEK293 cells. Two peaks corresponding to mNG-PIP4K dimers with either one or two mature fluorescent proteins were seen in the molecular brightness frequency distribution plot (**Figure 1E**). In contrast, the molecular brightness distributions measured for mNG-PIP5KA, mNG-PIP5KB, mNG-PIP5KC, mNG-zPIP5KA, and mNG-yMss4, all contain a single peak corresponding to the fluorescence of monomeric kinases labeled with a single fluorescent mNG (**Figure 1E**). Similar differences were observed when we compared the molecular brightness of recombinantly expressed and purified PIP4K and PIP5K that were chemically labeled with Alexa488 *in vitro* using Sortase mediated peptide ligation (**Figure 1–figure supplement 2A**).

Consistent with mNG-PIP4K and mNG-PIP5K binding to supported membranes with distinct oligomerization states, these two lipid kinases also exhibited strikingly different membrane diffusion coefficients. At low surface densities (~0.01 molecules/μm^2^) the mobility of mNG-PIP4K (0.04 μm^2^/sec) was much slower compared to mNG-PIP5K homologs and paralogs (0.15 – 0.18 μm^2^/sec) (**Figure 1F**). These diffusion coefficients were indistinguishable from those measured for kinases labeled with chemical dyes (**Figure 2C**), indicating that membrane binding of the mNG kinases in dilute cell lysate is indistinguishable from recombinantly purified and fluorescently labeled enzymes.

**Figure 2.**
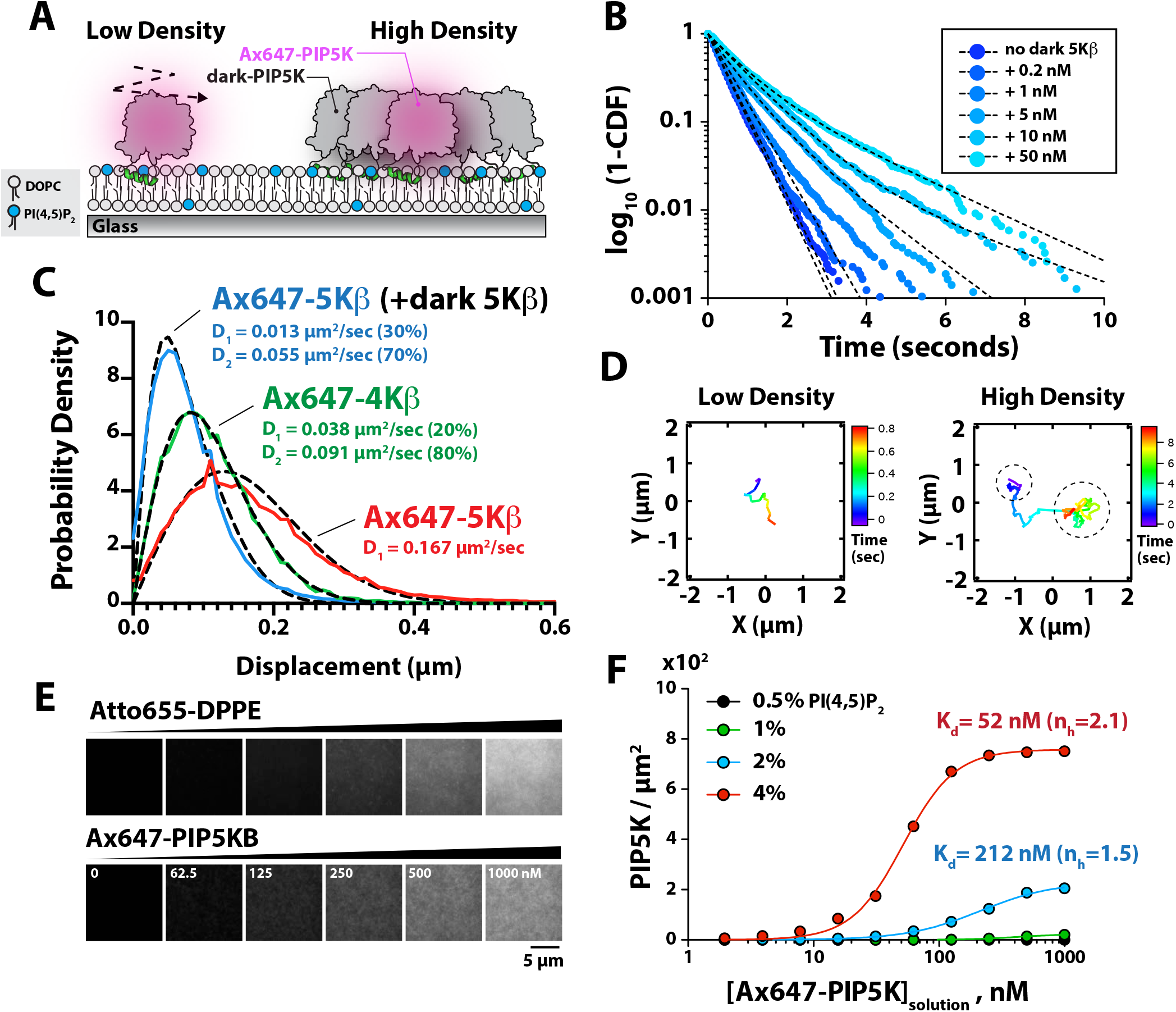
Protein density dependent changes in PIP5K membrane binding. **(A)** Supported lipid bilayer assay for characterizing Alexa647-PIP5KB single molecule membrane binding behavior on membranes containing low and high densities of PIP5KB. **(B)** Single molecule dwell times of Alexa647-PIP5KB measured in the presence of increasing concentrations of non-fluorescent PIP5KB. (C) Step size distribution measured in the presence of either 1 pM Alexa647-PIP5KB (red), 1 pM Alexa647-PIP5KB + 50 nM PIP5KB (blue), or 1 pM Alexa647-PIP4KB (green). Dashed black line represents the curve fit using the Stokes-Einstein equation (see methods). (D) Representative trajectories showing the time dependent movement of a single membrane bound Alexa647-PIP5KB molecule in the absence (low density’; left) or presence of 50 nM PIP5KB (‘high density’; right). **(B-D)** Membrane composition: 98% DOPC, 2% PI(4,5)P_2_. **(E)** Montage of images showing supported membranes with increasing densities of Atto655 lipids used to calibrate the molecular density of membrane bound Alexa647-PIP5KB. Membrane composition: 96% DOPC and 4% PI(4,5)P_2_. (F) Alexa647-PIP5KB binds cooperatively to membranes containing PI(4,5)P_2_ lipids. The density of membrane bound PIP5KB was measured in the presence of increasing solution concentrations of Alexa647-PIP5KB on membranes containing 0.5, 1, 2, or 4 % PI(4,5)P_2_ lipids.

### Membrane-mediated dimerization of PIP5K

The ability of PIP5K to associate with PI(4,5)P_2_ containing membranes as a monomer at low molecular densities (~0.01 PIP5K/μm^2^) confirms that PIP5K exists in a weak monomer-dimer equilibrium in solution. Membrane association reduces the effects of translational and rotational entropy, both of which oppose dimerization, enabling even weakly dimerizing species in solution to robustly dimerize on membranes (Chung et al. 2018; Lin et al. 2014). Thus we anticipated that membrane binding of PIP5K favors dimerization at some threshold membrane surface density. In order to characterize PIP5KB dimerization as a function of membrane surface density, we performed single molecule tracking experiments in the presence of low (~0.01 PIP5K/μm^2^) and high (~100 PIP5K/μm^2^) densities of membrane bound Ax647-PIP5KB (**Figure 2A**). These measurements reveal molecular binding dwell time on the membrane as well as diffusive mobility, both of which are affected by dimerization. In the presence of a low density of non-interacting Ax647-PIP5KB monomers, the distribution of single molecule dwell times could be fit to a single exponential with a characteristic dwell time of ~ 0.4 sec (**Figure 2B**). When we increased the solution concentration of PIP5KB we observed a second population of long dwelling Ax647-PIP5KB molecules, likely dimers or (possibly) higher order oligomers (**Figure 2B**). The resulting dwell time distributions were best fit to a two species model with two characteristic dwell times (**Figure 2B**). In addition to observing an enhancement in the dwell time, we also observed a protein surface density dependent decrease in the diffusion coefficient of membrane bound Ax647-PIP5KB (**Figure 2C**). Two-dimensional mobility on the membrane has previously been used as a highly effective measure of membrane surface dimerization reactions (Chung et al. 2018; 2019). A two species model was required to fit the step-size distribution of membrane bound Ax647-PIP5KB in the presence of high kinase density, which was similar to the step-size distribution of Ax647-PIP4KB measured at low molecular densities (**Figure 2C**). Examples of Ax647-PIP5KB molecules that transitioned between slow and fast diffusive states could also be seen when we inspected trajectories at an intermediate kinase density that favored PIP5K dimerization (**Figure 2D**).

We performed quantitative surface density measurements on supported membranes to assess how the membrane surface density of PIP5KB changes as a function of the PI(4,5)P_2_ lipid density and the kinase solution concentration. Using defined molar concentrations of Atto655-1,2-dipalmitoyl-sn-glycero-3-phosphoethanolamine (Atto655-DPPE) lipids incorporated into supported membranes, we calibrated the fluorescence intensity to measure the surface density of membrane bound Ax647-PIP5KB (**Figure 2E and Figure 2–figure supplement 1A**). These measurements revealed two non-linear membrane binding behaviors of Ax647-PIP5KB. First, the density of membrane bound Ax647-PIP5KB dramatically increased as a function of the PI(4,5)P_2_ density (**Figure 2F**). Second, increasing the solution concentration promoted cooperative membrane binding of Ax647-PIP5KB based on elevated protein densities (**Figure 2F)**. Fitting the membrane binding curves with a concerted model for cooperativity yielded dissociation constants of 212 nM and 52 nM for Ax647-PIP5KB in the presence of 2% and 4% PI(4,5)P_2_ lipids, respectively (**Figure 2F**). To determine if the density dependent change in the dwell time and the step size distributions of Ax647-PIP5KB were dependent on dimerization of the kinase domain, we sought to create a dimer interface mutant.

Inspection of the zPIP5K crystal structure (Hu et al. 2015) and primary amino acid sequence alignment revealed a high degree of conservation between PIP5K homologs (**Figure 3A-3B**). Based on conservation of the primary amino acid sequence, we mutated the dimer interface of human PIP5KB. Using single molecule TIRF-M, we compared the dwell times and diffusion coefficients of Ax647-PIP5KB and Ax647-PIP5KB (D51R) using low and high protein densities on SLBs. When measured at a low protein surface density (~0.01 molecule/μm^2^), the D51R mutant bound cooperatively to PI(4,5)P_2_ lipids in a manner that was indistinguishable from the wild-type kinase (**Figure 3C-3D**). Under these conditions, the molecular brightness distribution and diffusion coefficients of membrane bound Ax647-PIP5KB and Ax647-PIP5KB (D51R) were also similar (**Figure 3–figure supplement 1A-1B**). In contrast, single molecule membrane binding experiments executed using a high protein surface density (~100 molecules/μm^2^), revealed distinct membrane binding behaviors comparing Ax647-PIP5KB and Ax647-PIP5KB (D51R). The single molecule dwell time of Ax647-PIP5KB increased, while the dwell time of the D51R mutant remained unchanged in the presence of 50 nM unlabeled PIP5KB (**Figure 3E**). In addition, the diffusion coefficient of wild-type Ax647-PIP5KB decreased due to membrane mediated dimerization, while diffusivity of Ax647-PIP5KB (D51R) remained unchanged (**Figure 3F**).

**Figure 3.**
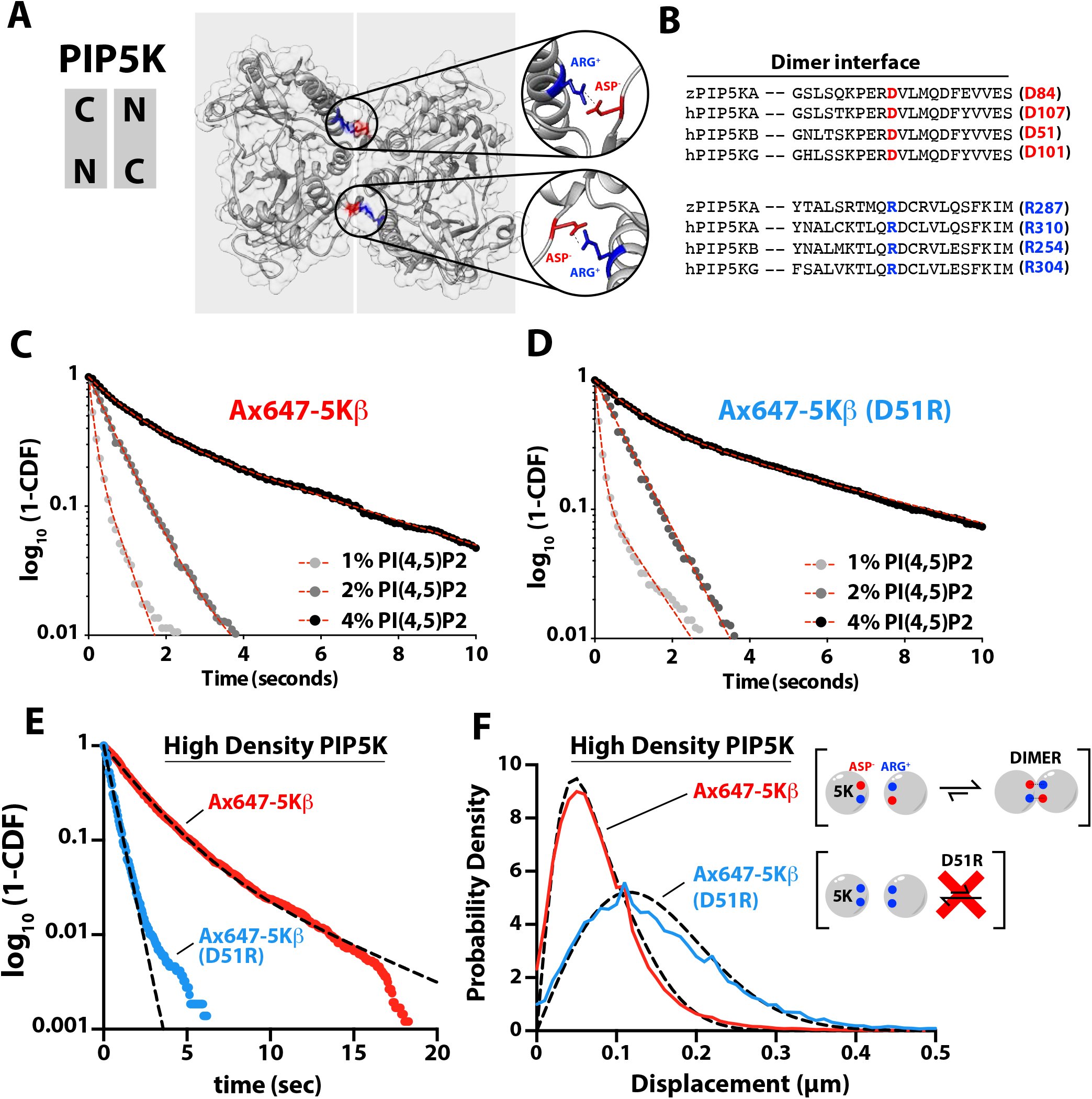
PIP5K binds cooperatively to PI(4,5)P_2_ independent of dimerization. **(A)** Kinase domain orientation in the zPIP5KA homodimer (4TZ7.pdb). Salt bridges formed between subunits are highlighted. (B) Sequence alignment between zPIP5K, hPIP5KA, hPIP5KB, hPIP5KG showing conservation of the dimer interface residues. **(C-D)** Dimerization is not required for cooperative PI(4,5)P_2_ binding. Single molecule dwell time distributions measured in the presence of (C) wild-type Alexa647-PIP5KB and (D) Alexa647-PIP5KB (D51R) on membranes containing different 1,2, or 4% PI(4,5)P_2_. (E-F) Single molecule dwell times and step size distributions measured in the presence of 50 nM PIP5KB on membranes containing 4% PI(4,5)P_2_ and 96% DOPC. (E) High density of PIP5KB increases the dwell time of Alexa647-PIP5KB, but not Alexa647-PIP5KB (D51R). (F) Membrane-mediated dimerization is responsible for the protein density dependent decrease in Alexa647-PIP5KB diffusion coefficient.

### Dimerization enhances PIP5K activity independently of increasing membrane recruitment

Previously, dimerization of zPIP5KA was shown to be essential for lipid kinase activity *in vitro* (Hu et al. 2015). Based on our single molecule dwell time analysis, dimerization of PIP5K is not required for cooperative binding to PI(4,5)P_2_ lipids (**Figure 3C-3D**). Comparing the kinetics of PI(4)P phosphorylation on SLBs, however, revealed differences in activity between the wild-type and the D51R mutant (**Figure 4A and Figure 4–figure supplement 1A-1B**). To determine if the differences in catalytic activity of the dimer mutant were caused by a reduction in membrane recruitment or dimerization induced change in catalytic efficiency, we simultaneously measured the kinetics of lipid phosphorylation and quantified the absolute density of membrane bound mNG-PIP5K. For both wild-type and D51R, membrane binding increased as a function of the PI(4,5)P_2_ membrane density (**Figure 4B**), with membrane binding of wild-type approximately 10-fold stronger compared to D51R. This was consistent with the enhanced dwell time mediated by dimerization of the kinase domain. To determine whether dimerization enhances the catalytic efficiency of PIP5K, we calculated the effective phosphorylation rate constants based on the calibrated surface density of mNG-PIP5KB (**Figure 4–figure supplement 2A-2C**). Compared to typical solution Michaelis-Menten kinetics, the density of membrane bound mNG-PIP5KB changes over the course of the lipid phosphorylation reactions described here. To account for this change, the effective phosphorylation rate per membrane bound PIP5KB molecule, *v_molecule_*(*t*), was calculated using the following equation:

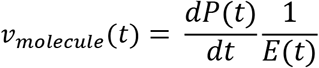

where *E* is the surface density of mNG-PIP5KB, and *P* is the surface density of PI(4,5)P_2_ on the membrane at any time in the reaction. The PI(4)P density at each time point, *S*(*t*), was approximated by subtracting *P*(*t*) from the total PIP lipid density, S_0_:

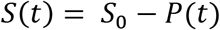

**Figure 4.**
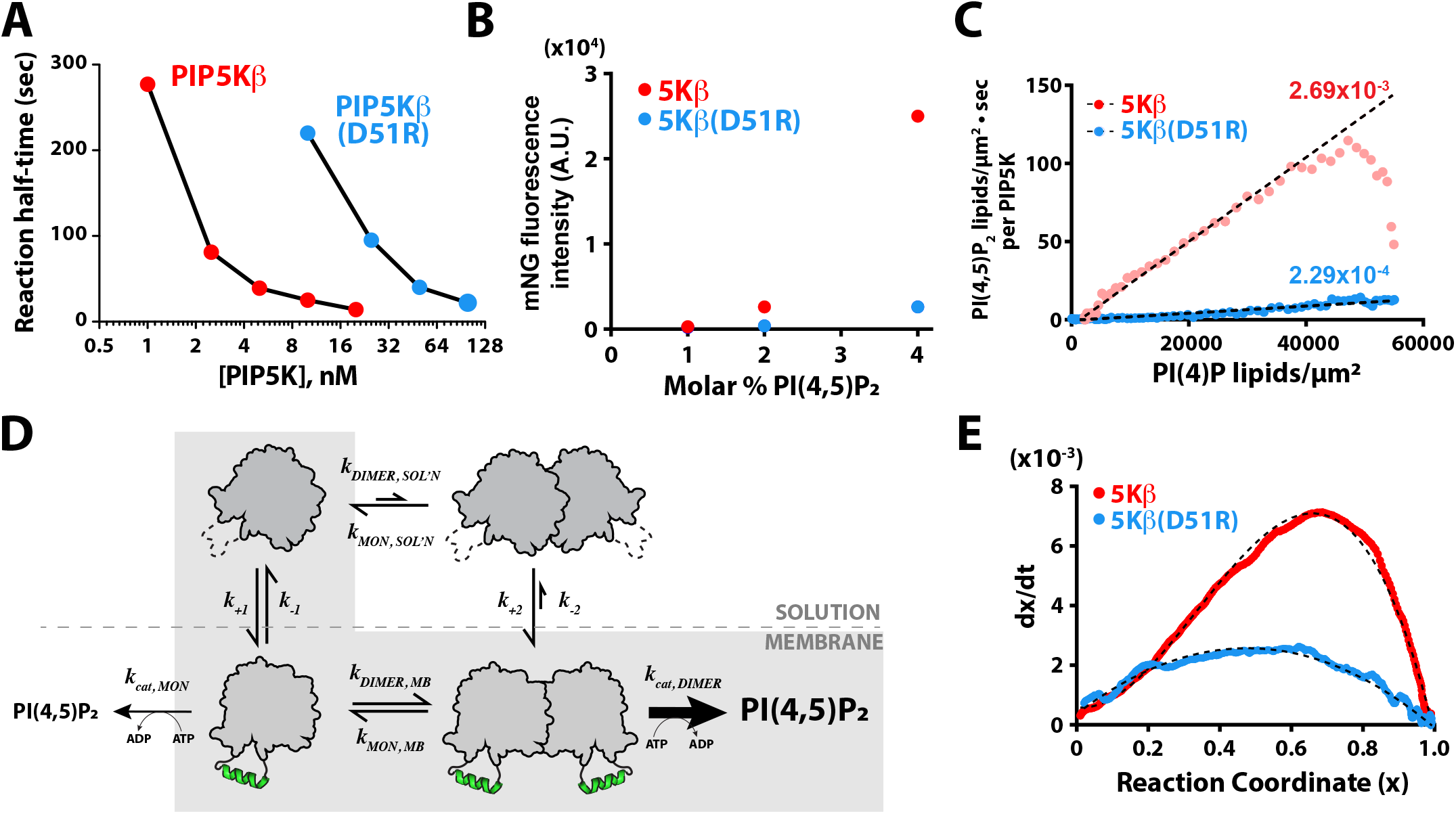
Membrane-mediated dimerization potentiates PIP5K lipid kinase activity. **(A)** Production of PI(4,5)P_2_ measured in the presence of varying concentrations of PIP5KB and PIP5KB (D51R). Initial membrane composition: 98% DOPC, 2% PI(4)P. (B) Equilibrium fluorescence intensity of membrane bound mNG-PIP5KB and mNG-PIP5KB (D51R) measured on membranes with increasing densities of PI(4,5)P_2_ lipids. (C) Membrane-mediated dimerization enhances PIP5K catalytic efficiency independent of membrane localization. Membrane localization mNG-PIP5K (wild-type and D51R) and production of PI(4,5)P_2_ were monitored simultaneously on supported membranes. Plot shows the number of PI(4,5)P_2_ lipids generated per μm^2^ per second per enzymes as a function of the substrate density. (D) Equilibrium diagram showing the mechanisms of PIP5K membrane binding and change in catalytic efficiency. (E) Feedback profiles for wild-type and D51R mutant PIP5KB. The following equations were used for curve fitting: 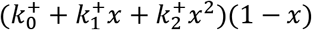 for PIP5KB and 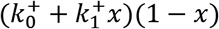 for PIP5KB (D51R). 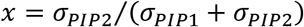.

By plotting *v_molecule_*(*t*) against the substrate density, *S*(*t*), we obtained a Michaelis-Menten plot with a slope equal to the effective per-molecule phosphorylation rate constant (**Figure 4C**). The calculated phosphorylation rate constant for the dimer mutant, PIP5KB (D51R), had a single rate constant of 2.3×10^-4^ lipids/μm^2^•sec per enzyme. Conversely, the phosphorylation rate constant for wild-type PIP5KB began with a slow rate and then transitioned to a rate of 2.7×10^-3^ lipids/μm^2^•sec per kinase as the reaction progressed. This acceleration in per-molecule reaction kinetics was consistent with a dimerization dependent increase in enzyme activity (**Figure 4D**), and establishes a positive feedback mechanism.

Comparing the shape of the kinetic traces for PIP5KB and PIP5KB (D51R) also revealed striking differences in the complexity of their positive feedback loops. To analyze the feedback profile of PIP5KB, we defined the reaction coordinate, x, as *x* ≡ *σPIP*2/(*σPIP*2 + *σPIP*1). The reaction rate on the coordinate can be expressed as:

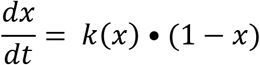

where *k*(*x*) is a function that characterizes the reaction rate, including any dependence on system composition (*x*), and can be expressed as a power series, *k*(*x*) = *k*_0_ + *k*_1_*x* + *k*_2_*x*^2^ +…. This provides a convenient way of examining the type of feedback; the order of feedback is revealed by the *x*-dependence of *k*(*x*). When the derivative of the PIP5K kinase reaction traces were plotted against the reaction coordinate, *x*, the curve displayed a high degree of asymmetry that required a second-order term to fit (*k_WT_*(*x*) = *k*_0_ + *k*_1_*x* + *k*_2_*x*^2^) (**Figure 4E**). In contrast, the *dx/dt* curve for the dimer mutant was parabolic and could be fit using an equation that describes an enzyme with first order positive feedback (*k*_*D*51*R*_(*x*) = *k*_0_ + *k*_1_*x*) based on product binding (**Figure 4E**). In previous studies, PIP5KB was shown to exhibit higher-order positive feedback (S. D. Hansen et al. 2019). One potential source of higher order positive feedback is through the ability of PIP5K to bind to multiple PI(4,5)P_2_ lipids. However, mapping the feedback strength based on the reaction coordinate revealed that dimerization is predominantly responsible for PIP5K higher-order positive feedback. Together, these results provide strong evidence that dimerization enhances the activity of PIP5K by both enhancing the membrane avidity and potentiating lipid kinase activity through a mechanism that is consistent with allosteric regulation.

### PIP5K dimerization increases the stability of PIP compositional patterns

The PIP5KB positive feedback mechanism previously enabled us to reconstitute a bistable lipid kinase-phosphatase competitive reaction that breaks symmetry and produces PI(4)P and PI(4,5)P_2_ compositional patterns on supported membranes (**Figure 5A**) (S. D. Hansen et al. 2019). Based on the ability of PIP5KB to undergo membrane mediated dimerization, and the corresponding nonlinear positive feedback, we hypothesized the kinetic bistability in this system may be enhanced by dimerization of PIP5K. To test this hypothesis, we compared the ability of PIP5KB and the D51R dimer mutant to form bistable PIP compositional patterns in the presence of an opposing 5’-phosphatase. In the presence of 50 nM PIP5KB (WT or D51R) and 30 nM DrrA-OCRL, we found that the activity of the D51R dimer mutant was strongly impaired compared to wild-type PIP5KB (**Figure 5B**). Restoring the PIP5KB dimer interface with a charge reversal mutation, D51R/R254D, allowed us to reconstitute bistable compositional patterns that were indistinguishable from those formed in the presence of wild-type PIP5K (**Figure 5B and Figure 4–figure supplement 1C**). Taking into consideration the weakened positive feedback of the dimer mutant, we raised the solution concentration of PIP5KB (D51R) until we were able to balance the opposing 5’-phosphatase activity. Using a 20-fold higher concentration of PIP5KB (D51R), compared to wild-type, we identified a concentration regime that allowed us to reconstitute PIP compositional patterns (**Figure 5B**). During the early stages of pattern formation, the surface area and morphology looked very similar to compositional patterns reconstituted in the presence of wild-type PIP5KB. However, several minutes after observing symmetry breaking of the PIP lipids the PI(4,5)P_2_ compositional patterns generated by PIP5KB (D51R) were consumed by the surrounding 5’-phosphatase dominated reaction (**Figure 5B**).

**Figure 5.**
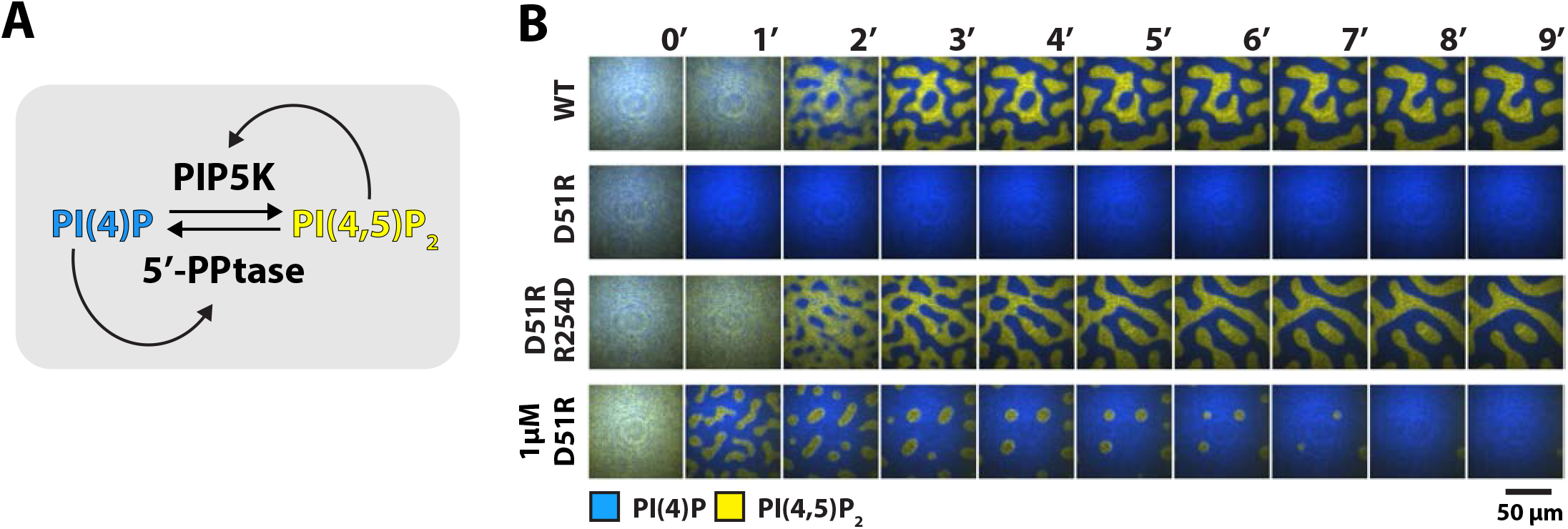
PIP5K dimerization stabilizes PIP compositional patterns. **(A)** Schematic of competitive reaction containing PIP5K and 5’-PPtase. **(B)** Representative image sequences showing the reconstitution and visualization of bistable PIP compositional patterns in the presence 50 nM PIP5KB (wild-type, D51R, or D51R/R254D), 30 nM DrrA-OCRL, 20 nM Alexa488-PLCδ, 20 nM Alexa647-DrrA. Compositional patterns were formed in the presence of 1 μM PIP5KB (D51R). Initial membrane composition: 96% DOPC, 2% PI(4)P, 2% PI(4,5)P_2_.

### Reaction trajectory variation is enhanced by membrane-mediated dimerization

We previously reported that PIP5K dependent lipid phosphorylation reactions exhibit a high degree of reaction trajectory variation when reconstituted on supported lipid bilayers that are partitioned into micron length scale membrane corrals (S. D. Hansen et al. 2019). Based on the coupling induced by membrane-mediated dimerization, we hypothesized that the dimerization could provide the molecular basis for the previously observed enhanced reaction trajectory variation. To measure dimerization dependent differences in reaction trajectory variation we microfabricated an array of 5 μm x 5 μm chromium barrier onto the underlying glass coverslip. This approach allowed us to visualize hundreds of identical membrane reactions in parallel that continuously exchange with the surrounding solution environment. Under these conditions, PIP5KB reaction trajectories displayed a high degree of kinetic heterogeneity (**Figure 6A-6B**). While the time to finish was highly heterogeneous, the reaction rate of the initial and later part of the reaction was quite homogeneous. This was most noticeable when plotting the kinetic data in the reaction coordinate plot (**Figure 6B**). In contrast, reactions reconstituted in the presence of PIP5KB (D51R) showed little heterogeneity confirming that dimerization strongly enhanced variation in corral reaction trajectory (**Figure 6B**). Overall, the observed reaction heterogeneity was primarily driven at higher product concentrations, where higher-order positive feedback dominates. Formation of the PIP5K dimer creates a highly active kinase with a strengthened positive feedback and faster reaction velocity.

**Figure 6.**
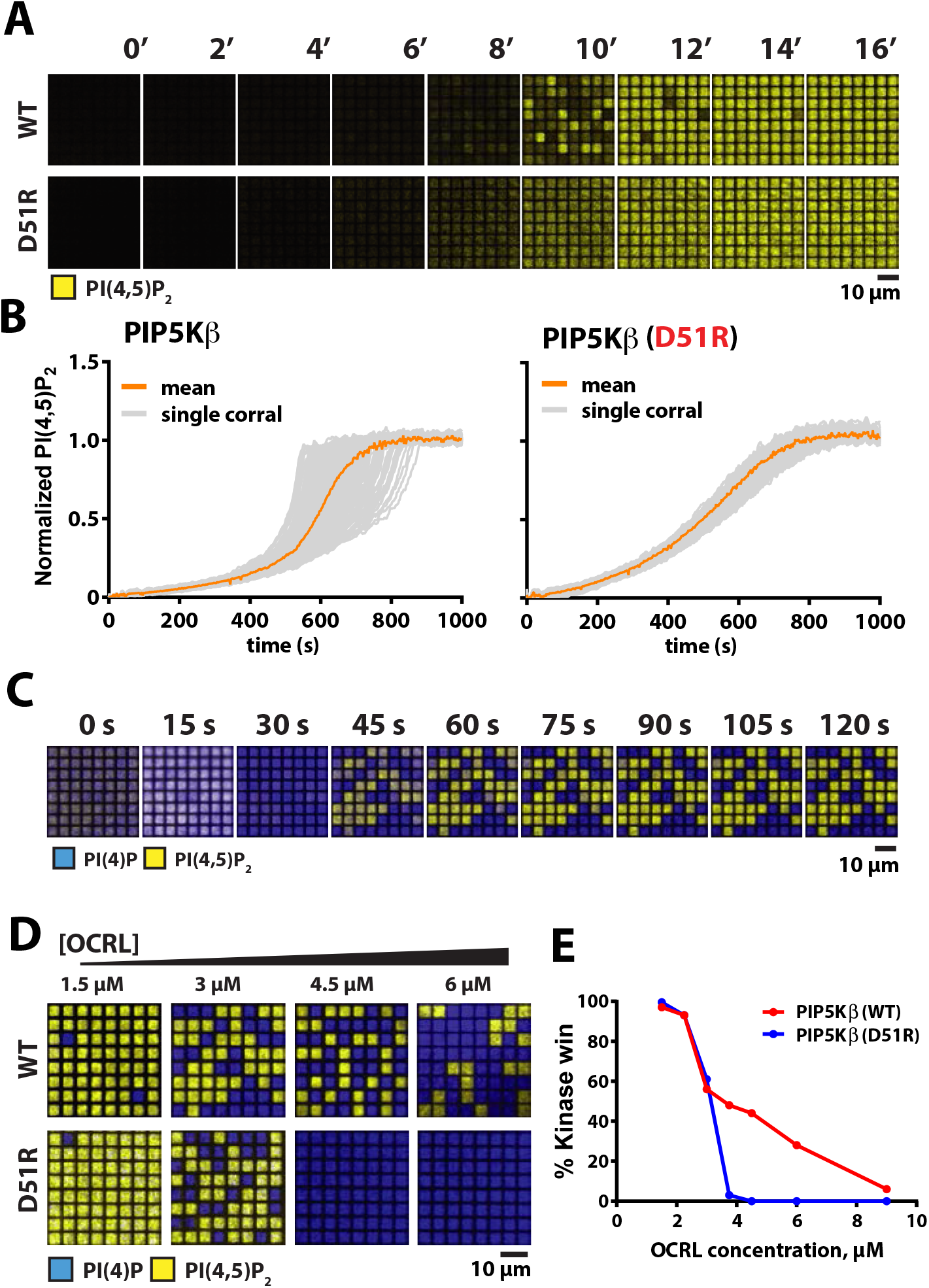
Reaction trajectory variation based on stochastic mem brane-mediated dimerization. **(A)** Lipid phosphorylation reactions reconstituted in 5 μm x 5 μm chromium patterned supported membranes in the presence of 5 nM PIP5K and 10 nM PIP5K (D51R). Reactions were visualized using 20 nM Alexa488-PLCδ. Initial membrane composition: 96% DOPC, 4% PI(4)P. (B) Reaction trajectories plots from (A). (C) Time sequence of bistable kinase-phosphatase reaction reconstituted in 5μm x 5μm corrals in the presence of 50 nM PIP5K, 30 nM DrrA-OCRL, 20 nM Alexa488-PLCδ, and 20 nM Alexa647-DrrA. **(D)** Dimerization enhances ability of PIP5K to win competitive bistable kinase-phosphatase reaction in the presence of OCRL. Reactions were performed in the presence of 3 μM OCRL and either 18 nM PIP5KB or 370 nM PIP5KB (D51R). (E) Quantification of final reaction outcome in (D). **(C-E)** Initial membrane composition: 96% DOPC, 2% PI(4)P, and 2% PI(4,5)P_2_.

### Stochastic effects can enhance bistability

Nonlinear positive feedback is sufficient to establish bistability in a competitive reaction, even on scales over which stochastic variations average to essentially zero. The stable patterns seen in **Figure 5B** (WT and D51R/R254D) are likely a representation of this non-stochastic bistability, with two intrinsically stable steady states. The dimer mutant, however, shows only linear positive feedback (see **Figure 4E**) but still shows at least transient bistability in **Figure 5B**. This is a manifestation of stochastic bistability, referring to bistable behavior in systems that inherently lack two stable steady states (Bishop and Qian 2010; S. D. Hansen et al. 2019; Artyomov et al. 2007; To and Maheshri 2010). For the WT PIP5K, dimerization enhances stochastic variation in reaction rate and thus we hypothesize will expand the range of conditions over which bistable behavior is observed, even spanning beyond the boundaries of intrinsic bistability. To test this hypothesis, we examine competitive reactions with OCRL phosphatase for WT and D51R under geometric confinement, where stochastic effects are prominent. Reconstitution of the WT PIP5K and OCRL competitive reactions in 5 μm x 5 μm membrane corrals reveals strong bistability, reaching final membrane compositions in each corral consisting of steady-states dominated by either PI(4)P or PI(4,5)P_2_ composition (**Figure 6C**). To compare with the dimer mutant, we titrate the competing OCRL concentration against fixed concentrations of either PIP5KB and PIP5KB (D51R) and examine the resultant bistability (**Figure 6D**). WT kinase exhibits a much more robust bistability, which spans a substantially wider range of competing phosphatase concentrations than observed for the dimer mutant. Quantification of this effect is plotted in **Figure 6E**. Dimerization of PIP5K thus ensures that the kinase–phosphatase competitive reactions can achieve a bistable response over a broad range of opposing 5’-phosphatase activity.

## DISCUSSION

### Dimerization as a mechanism for allosteric control of PIP5K activity

Cooperative PI(4,5)P_2_ binding and membrane-mediated dimerization provide synergistic mechanisms to increase the rate of PI(4,5)P_2_ production through the enhanced localization and increased catalytic efficiency of PIP5K (**Figure 7**). Supporting a mechanism of allosteric regulation, our results indicate that dimerization enhances PIP5K lipid kinase activity by directly increasing *k_cat_*. This effect is independent of increasing membrane avidity of PIP5K. Due to a lack of structural biochemistry, there is currently a gap in a knowledge concerning the role dimerization, PIP lipid binding, and the nucleotide state serve in regulating conformational states of PIP5K. Given our limited structural understanding of PIP5K (Liu et al. 2016; Hu et al. 2015; Muftuoglu et al. 2016), some researchers have used molecular dynamic simulations to elucidate how membrane docking of PIP5K is controlled (Amos et al. 2019). Working under the assumption that PIP5K is constitutively dimeric, Amos et al. reported that only a single kinase domain can engage substrate when the dimer is docked on a PI(4)P containing membrane. If this mechanism is accurate, membrane-mediated dimerization is expected to enhance processivity of membrane bound PIP5K by allowing kinase domains to toggle between states of catalysis and membrane binding. This mechanism would ensure that dimeric PIP5K remains bound to the membrane during enzyme catalysis, while the monomeric PIP5K would likely catalyze a single PI(4)P phosphorylation reaction before dissociating. According to our single molecule biophysical studies, we believe that both subunits of the PIP5K kinase domain can engage the membrane. This is based on the enhanced dwell time and slower diffusion coefficient observed when Ax647-PIP5K is visualized is high protein density (i.e. ~100 molecules/μm^2^). Incorporating reversible dimerization, catalysis, and PI(4,5)P_2_ binding into future molecular dynamic simulations could provide new insight about the mechanism of membrane docking.

**Figure 7.**
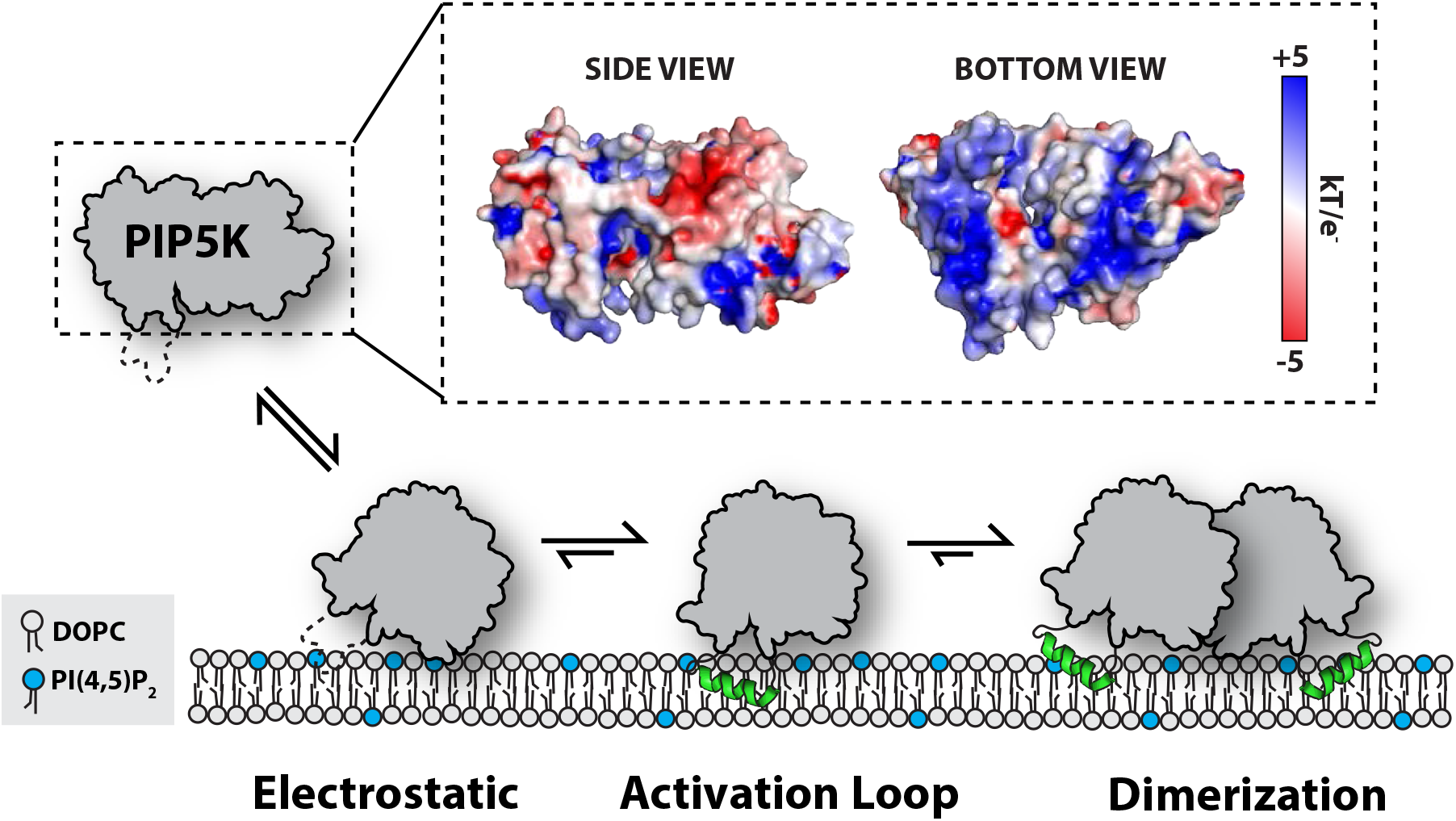
Model for PIP5K membrane-mediated dimerization. Mediated by electrostatic interactions, PIP5K can associate with membranes as a monomer. Insertion of the activation loop stabilizes membrane association and enables PIP5K to catalyzes the phosphorylation of PI(4)P to generate PI(4,5)P_2_. Increasing densities of PI(4,5)P_2_ and membrane bound PIP5K promotes membrane-mediated dimerization, which leads to enhanced catalytic efficiency.

### PIP5K dimerization increases reaction trajectory variation

Human proteomic data has estimated the cellular concentration of PIP5K to be 10-20 nM in mammalian cells (Hein et al. 2015). Based on the weak monomer-dimer equilibrium, PIP5K is predicted to exist predominantly as a monomer in the cytoplasm. Cooperative PI(4,5)P_2_ binding, membrane-mediated dimerization, and low molecular copy number of PIP5K provides several mechanisms for enhancing stochasticity in membrane associated lipid phosphorylation reactions. Previously, we reported that reaction velocity fluctuations driven by stochastic binding and unbinding of PIP5K on the membrane could drive the system to a predominantly PI(4,5)P_2_ state, even in the presence of high levels of opposing lipid phosphatase activity that were sufficient to drive the system predominantly to the PI(4)P state in bulk (S. D. Hansen et al. 2019). The resulting emergent property is that the reaction outcome depends on system size, which we termed stochastic geometry sensing; related scale-dependent phenomena have been called reaction inversion in some theoretical work (Ramaswamy et al. 2012). Here we report that dimerization is responsible for the higher order positive feedback exhibited by PIP5K, and this contributes to the robustness with which this system exhibits stochastic geometry sensing. Without dimerization, PIP5K displays a simple linear positive feedback profile. Comparing the lipid phosphorylation reaction trajectory variation of PIP5K and PIP5K (D51R) highlights how dimerization enhances stochasticity of lipid kinase phosphorylation reactions and will thus amplify any effects dependent on stochastic fluctuations. Given the broad dynamic range of kinase densities and activities that can emerge from these biochemical mechanisms, stochasticity in PIP5K signaling has the potential to strongly influence signaling events orchestrated at the plasma membrane in living cells.

### PIP5K localization and lipid kinase activity in cells

The strong lipid phosphorylation activity of PIP5K observed *in vitro* raises questions about the steady-state localization and activity of PIP5K in cells. Although new molecular mechanisms concerning PIP5K activation have been revealed through single molecule characterization of PIP5K *in vitro*, it remains challenging to interpret how dimerization, PI(4,5)P_2_ binding, and interactions with peripheral membrane proteins regulate membrane localization of PIP5K *in vivo*. Complicating researcher interpretation of PIP5K localization and dynamics, PIP5K paralogs have the potential to heterodimerize (Lacalle et al. 2007). Determining how cells regulate the strength and duration of PIP5K lipid phosphorylation reactions during receptor activation will help elucidate the role cooperative PI(4,5)P_2_ binding and dimerization serve in enhancing PIP5K dependent production of PI(4,5)P_2_ during cell signaling. Receptor activation could shift the steady-state localization of PIP5K to favor membrane dependent dimerization and enhanced lipid kinase activity. This could be achieved by increasing PIP5K membrane recruitment through interactions with receptors, endocytic machinery, or small GTPases (Halstead et al. 2010; Honda et al. 1999; Funakoshi, Hasegawa, and Kanaho 2010). If the steady-state localization of PIP5K exists near a threshold for activation, a minor enhancement in localization could promote PIP5K dimerization and trigger the PI(4,5)P_2_ dependent positive feedback loop. Future biochemical characterization of PIP5K will enable researchers to generate better separation of function mutants that allow the functional significance of specific protein-protein and protein-lipid interactions to be studied in cell signaling.

One challenge in establishing an accurate model for PIP5K membrane binding and catalysis is the gap in knowledge concerning how PIP5K binds to PI(4,5)P_2_ lipids. Previously published cell biology data reported that several basic residues are critical for PIP binding and plasma membrane localization in cells (Fairn et al. 2009). However, these basic residues are dispensable for PIP5K membrane binding and activity *in vitro* (S. D. Hansen et al. 2019). Looking at the crystal structure of zPIP5KA (Hu et al. 2015; Muftuoglu et al. 2016) there are several motifs of unresolved electron density and domains extending beyond the N- and C-terminus of the kinase domain for which we lack high resolution structural data. Defining the functional relevance of those domains in controlling PIP5K membrane association could provide new insight about the molecular basis of PI(4,5)P_2_ binding specificity. By determining the molecular basis of the PIP5K and PI(4,5)P_2_ interaction cell biologists can instead use fluorescently labeled PIP5K to directly visualize local fluctuations in PI(4,5)P_2_ inferred from changes in the single molecule dwell time of membrane bound PIP5K, such as has been done with Grb2 and other signaling molecules to monitory receptor activity (Chen et al. 2021; 2018). We already know from previous work that PIP5K becomes enriched at sites of clathrin-mediated endocytosis (Nakano-Kobayashi et al. 2007) and focal adhesions (Ling et al. 2002; Di Paolo et al. 2002). It remains unclear whether localization of PIP5K to these cellular structure requires interactions with PI(4,5)P_2_ lipids, dimerization of the kinase domain, or recruitment by peripheral membrane proteins.

### Negative regulation of PIP5K signaling

The strong positive feedback loop displayed by PIP5K raises many questions about how lipid kinase activity is turned off once production of PI(4,5)P_2_ ramps up. Left unregulated, the PIP5K positive feedback loop has the potential to generate excessively high concentrations of PI(4,5)P_2_ in cells. This would be detrimental to numerous signaling pathways which rely on cellular PIP lipid homeostasis. By limiting the concentration of freely available PI(4,5)P_2_ cells can potentially control the strength and duration of the PIP5K positive feedback loops. *In vitro*, PIP5K interacts strongly with supported membranes containing 2-4% PI(4,5)P_2_. Although the total concentration of PI(4,5)P_2_ is estimated to be 0.5 – 5% percent in the plasma membrane (Wenk et al. 2003; Mitchell, Ferrell, and Huestis 1986; Nasuhoglu et al. 2002), the concentration of freely available PI(4,5)P_2_ is likely lower. PI(4,5)P_2_ sequestering proteins, like MARCKS, are present at micromolar concentrations in cells (Brudvig and Weimer 2015). As a result, there is likely a lower concentration of freely available PI(4,5)P_2_ at the plasma membrane. Lipid phosphatases, phosphatidylinositol-3 kinase (PI3K), and phospholipases also regulate the conversion of PI(4,5)P_2_ into other lipid species and secondary metabolites (Balla 2013). In the case of early phagocytosis, macrophages can put the brakes on PIP5K activity by activating phospholipase C, which cleaves the inositol head group from PI(4,5)P_2_ to produce second messenger, IP3 and DAG (Rhee 2001). Parallel to this pathway, PI3K phosphorylates PI(4,5)P_2_ to generate PI(3,4,5)P_3_. Both enzymatic reactions reduce the amplitude and duration of PI(4,5)P_2_ spikes. By understanding how lipid kinases, phosphatases, and sequestering molecules regulate the activity of PIP5K will be critical for understanding how cells control PI(4,5)P_2_ lipid homeostasis. Considering the myriad of human diseases linked to the disruption of PIP lipid homeostasis, there is much to gain by understanding the molecular mechanisms that regulate the spatiotemporal dynamics of PIP5K lipid kinase activity in cells.

## Acknowledgments

We thank members of the Hansen lab for critical reading of our manuscript. Funding was provided by NIH grant P01 AI091580 (J.T.G), by the Novo Nordisk Foundation Challenge Programme as part of the Center for Geometrically Engineered Cellular Systems (J.T.G.), National Research Service Award Postdoctoral Fellowship (S.D.H., F32 GM111010-02), University of Oregon Startup funds (S.D.H.), and NSF CAREER Award (S.D.H., MCB-2048060).

## Author contributions

Biochemistry & Reagents, S.D.H. and A.A.L.; Experiments, S.D.H. and A.A.L.; Data Analysis, S.D.H. and A.A.L.; Conceptualization of data, S.D.H., A.A.L., and J.T.G.; Interpretation, S.D.H., A.A.L., and J.T.G.; Edited manuscript, S.D.H., A.A.L., and J.T.G.; wrote original manuscript, S.D.H..

## Data and materials availability

All the information needed for interpretation of the data is presented in the manuscript or the supplemental material. Plasmids and reagents related to this work are available upon request.

## SUPPLEMENTAL FIGURE LEGENDS

**Figure 1 – figure supplemental 1**

**mNeonGreen calibration curve for single molecule cell lysate assay**

**(A)** The fluorescence of emission of purified mNG was measured over a broad range of solution concentration using a BioTek 96-well plate reader. Representative standard curve from a single experiment was fit using standard linear regression.

**Figure 1 – figure supplemental 2 Molecular brightness analysis of PIP4K and PIP5K**

**(A)** Single molecule brightness distributions for Sortase labeled Alexa488-PIP4K and Alexa488-PIP5K bound to supported membranes containing 96% DOPC and 4% PI(4,5)P_2_.

**Figure 2 – figure supplemental 1**

**Calibration of Alexa647-PIP5KB membrane surface density measurements**

**(A)** Calculation of scaling factor used from comparing the fluorescence of Atto640-DPPE and Alexa647-PIP5KB. The fluorescence intensity of small unilamellar vesicles containing vary concentrations of Atto655-DPPE lipids or purified Alexa647-PIP5KB were measuring using wide-field fluorescence microscopy. Data points represent the average fluorescence intensity of 20 images acquired using the identical camera settings, objective, laser power, and filters.

**Figure 3 – figure supplemental 1**

**Dimer interface mutation does not alter diffusion of PIP5K at low protein densities**

**(A-B)** Step-size distributions are indistinguishable when measured in the presence of **(A)** 5 pM Alexa647-PIP5KB or **(B)** 5 pM Alexa647-PIP5KB (D51R). Membrane composition: 98% DOPC, 2% PI(4,5)P_2_.

**Figure 4 – figure supplemental 1**

**Fluorescence correlation spectroscopy membrane density calibration**

**(A)** Auto-correlation of his10-mNeonGreen (his10-mNG) measured by FCS. Membrane composition: 98% DOPC, 2% NiNTA-DGS. (**B-C**) Calibration of mNG-PIP5K and mNG-PIP5K (D51R) membrane surface density using his10-mNG control. Increasing concentration of respective mNG-PIP5K proteins were incubated with support membranes containing 96% DOPC and 4% PI(4,5)P_2_.

**Figure 4 – figure supplemental 2**

**Dimerization enhances PIP5KB lipid kinase activity**

**(A-C)** Kinetic traces measuring phosphorylation of PI(4)P by **(A)** PIP5K, **(B)** PIP5KB (D51R), and **(C)** PIP5KB (D51R/R254D). The production of PI(4,5)P_2_ was monitored by the presence of 20 nM Alexa488-PLCδ. Initial membrane composition: 98% DOPC, 2% PI(4)P.

## MATERIALS AND METHODS

### Molecular Biology

Genes coding for *L. pneomophila* DrrA/SidM (Accession #Q5ZSQ3.1), human phosphatidylinositol 4,5-bisphosphate phosphodiesterase delta-1 PH domain (PLCδ ccession #P51178.2), human oculocerebrorenal syndrome of Lowe inositol polyphosphate 5-phosphatase (OCRL; Uniprot #Q01968), and human phosphatidylinositol 4-phosphate 5-kinase type-1 beta (hPIP5KB; Uniprot #O14986) were derived from codon optimized genes synthesized by GeneArt (Invitrogen). The gene encoding yeast Mss4 (Uniprot #P38994) was obtained by PCR amplification from yeast genomic DNA (S288C, *Saccharomyces cerevisiae).* The gene encoding mouse PIP5KA isoform 1 was purchased as a cDNA clone from Horizon Discovery (cat# MMM1013-202762630, Uniprot #P70182). The gene encoding human PIP5KC1 (Uniprot #O60331) was provided by Peter Bieling (Max Planck Institute of Molecular Physiology/Dortmund, Germany). The plasmids containing codon optimized zebrafish PIP5KA (zPIP5KA 49-431aa, Accession # NP_001018438.1) for expression in bacteria was kindly provided by Jian Hu (Michigan State University, Department of Biochemistry and Molecular Biology). Gene sequences were subcloned into bacterial and baculovirus protein expression vectors containing coding sequences with different solubility and affinity tags. PIP5K genes were cloned into a modified FAST Bac1 vector using ligation independent cloning or Gibson assembly (Gibson et al. 2009). The complete open reading frame of all vectors used in this study were sequenced to ensure the plasmids lacked deleterious mutations.

### Purification of BACMID DNA

To create BACMID DNA, FASTBac1 plasmids were transformed into DH10 Bac cells and plated on agar containing 50 μg/mL kanamycin, 10 μg/mL tetracycline, 7 μg/mL gentamycin, 40 μg/mL X-GAL, and 40 μg/mL IPTG. After 2-3 days of growth at 37°C, positive clones were isolated based on blue-white colony selection. Single white colonies were picked and struck on a BACMID agar plate for a second round of selection. BACMIDs were purified from 3 mL bacterial cultures grown overnight in TPM. Bacteria are centrifuged and resuspended in 300 μL of buffer containing 50 mM Tris [pH 8.0], 10 mM EDTA, 100 μg/mL RNase A (Qiagen PI buffer). Bacteria were then lysed by adding 300 μL of buffer containing 200 mM NaOH, 1% SDS (Qiagen P_2_ buffer). Neutralize lysis buffer by adding 300 μL of 4.2 M Guanidine HCl, 0.9 M KOAc [pH 4.8] (Qiagen N3 buffer). Centrifuge sample at 23°C for 10 minutes at 14,000 x g. Remove supernatant and combine with 700 μL 100% isopropanol. Centrifuge sample at 23°C for 10 minutes at 14,000 x g. Remove supernatant and add 200 μL of 70% ethanol. Centrifuge sample at 23°C for 10 minutes at 14,000 x g. Remove supernatant and add 50 μL of 70% ethanol. Centrifuge sample at 23°C for 10 minutes at 14,000 x g. Remove supernatant and dry DNA pellet slightly in biosafety hood. Solubilize DNA with 40 μL of sterile water. Resuspend DNA pellet by tapping side of micro-centrifuge tube 15-20 times. Quantify concentration of DNA using NanoDrop (typically 200-300 ng/μL). Immediately used BACMID DNA for transfection of Sf9 cells. Remaining BACMID DNA can be stored in −20°C freezer.

### Baculovirus production

Baculoviruses was generated by transfecting 1 x 10^6^ *Spodoptera frugiperda* (Sf9) insect cells plated for 24 hours in a Corning 6-well plastic dish (Cat# 07-200-80) containing 2 mL of ESF 921 Serum-Free Insect Cell Culture media (Expression Systems, Cat# 96-001, Davis, CA.). All media contains 1x concentration of Antibiotic-Antimycotic (Gibco/Invitrogen, Cat#15240-062). For transfection, 5-7 μg BACMID DNA was incubated with 4 μL Fugene (Thermo Fisher, Cat# 10362100) in 200 μL of ESF 921 media for 30 minutes at 23°C. BACMID DNA and Fugene were added dropwise to 6-well dish. Media change before and after addition of transfection reagent is unnecessary. After 4-5 days of transfection, viral supernatant (termed ‘P0’) is harvested, centrifuged, and used to infect 7 x 10^6^ *Sf9* cells plated for 24 hours in 10 cm tissue culture grade petri dish containing 10 mL of ESF 921 media and 10% Fetal Bovine serum (Seradigm, Cat# 1500-500, Lot# 176B14). After 4 days, viral supernatant (termed ‘P1’) is harvested and centrifuged to remove cell debri. Typical P1 viral titer yield is 10-12 mL. The P1 viral titer is expanded in 100 mL *Sf9* cell culture grown to a density of 1.25-1.5 x 10^6^ cells/mL in a sterile 250 mL polycarbonate Erlenmeyer flask with vented cap (Corning, #431144). We typically transduce 100 mL Sf9 culture with a concentration of 1% vol/vol of PI viral titer. Remaining PI virus is frozen as 1.5 mL aliquots that are stored in the −80°C freezer. The 10% Fetal Bovine serum serves as a cryo-protectant. After 4 days, viral supernatant (termed ‘P2’) is harvested, centrifuged, and 0.22 μm filtered in 150 mL filter-top bottle (Corning, polyethersulfone (PES), Cat#431153). The P2 viral supernatant is used for protein expression in High 5 cells grown in ESF 921 Serum-Free Insect Cell Culture media. The MOI for optimal protein expression is determined empirically to minimize cell death and maximize protein yield (typically 1.5-2% vol/vol final concentration of P2 virus).

#### Protein purification

##### PIP5KA, PIP5KB, PIP5KC, Mss4

Gene sequences encoding PIP5K family proteins were cloned into a FastBac1 vector in frame with a N-terminal his6-MBP-(Asn)_10_-TEV-GGGGG. ES-Sf9 cells were infected with baculovirus using an optimized multiplicity of infection (MOI), typically 1.5–2% vol/vol, was determined empirically from small-scale test expression (25-50 mL culture). Infected cells were typically grown for 48 hours at 27°C in ESF 921 Serum-Free Insect Cell Culture medium (Expression Systems, Cat# 96-001-01). Cells are harvested by centrifugation, washed with 1x PBS [pH 7.2], flash frozen in 50 mL tubes using liquid nitrogen and then stored in the −80°C freezer. For purification, frozen cells were thaw in an ambient water bath and lysed into buffer containing 50 mM Na_2_HPO_4_ [pH 8.0], 10 mM imidazole, 400 mM NaCl, 1 mM PMSF, 5 mM BME, 100 μg/mL DNase, 1 Sigma protease inhibitor cocktail EDTA free per 100 mL lysis buffer using a dounce homogenizer. Lysate was centrifuged at 16,000 rpm (35,172 x *g*) for 60 minutes in a Beckman JA-17 rotor at 4°C. Lysate was then batch bound to 5 mL of Ni-NTA Agarose (Qiagen, Cat# 30230) resin at 4°C for 1 hour in a beaker set on a stir plate. Resin was then collected in 50 mL tubes, centrifuged, and washed with buffer containing 50 mM Na_2_HPO_4_ [pH 8.0], 10 mM imidazole, 400 mM NaCl, and 5 mM BME before being transferred to gravity flow column. NiNTA resin with his6-MBP-(Asn)_10_-TEV-GGGGG-PIP5K (or Mss4) was then washed with 100 mL of 50 mM Na_2_HPO_4_ [pH 8.0], 30 mM imidazole, 400 mM NaCl, and 5 mM BME buffer and then eluted into buffer containing 500 mM imidazole. Peak fractions were pooled, combined with 200 μg/mL his6-TEV(S291V) protease, and dialyzed against 4 liters of buffer containing 20 mM Tris [pH 8.0], 200 mM NaCl, 2.5 mM BME for 16-18 hours at 4°C. Dialysate was then combined 1:1 with 20 mM Tris [pH 8.0], 1 mM DTT (~100 mM NaCl final). Precipitation was removed by centrifugation and 0.22 μm syringe filtration. Clarified dialysate was then bound to a MonoS cation exchange column (GE Healthcare, Cat# 17-5168-01) equilibrated in 20 mM Tris [pH 8.0], 100 mM NaCl, 1 mM TCEP buffer. Proteins were resolved over a 10-100% linear gradient (0.1-1 M NaCl, 45 CV, 45 mL total, 1 mL/min flow rate). PIP5K homologs and paralogs typically eluted from the MonoS column in the presence of 370-450 mM NaCl. Peak fractions containing PIP5K (or Mss4) were pooled, concentrated in a 30 kDa MWCO Vivaspin 6 centrifuge tube (GE Healthcare, Cat# 28-9323-17), and loaded onto a 24 mL Superdex 200 10/300 GL (GE Healthcare, Cat# 17-5174-01) size exclusion column equilibrated in 20 mM Tris [pH 8.0], 200 mM NaCl, 10% glycerol, 1 mM TCEP. Peak fractions were concentrated in a 30 kDa MWCO Vivaspin 6 centrifuge tube and snap frozen at a final concentration of 10-20 μM using liquid nitrogen.

##### zPIP5K

The coding sequence of zebrafish PIP5KA (49-431aa) was expressed in *E. coli* Rosetta2 pLysS as a C-terminal his6 fusion protein. Bacteria were grown at 37°C in LB until OD_600_=0.5. Due to poor solubility of zPIP5KA, we used 8-10 liters of total cell culture volume. Cultures were shifted to 23°C for approximately a 1 hour, induced with 0.1 mM IPTG, and allowed to express protein for 16 hours before being harvested. Cell harvested by centrifugation were resuspended in lysis buffer made of 50 mM HEPES, pH 7.3, 300 mM NaCl, 5% Glycerol, 0.5% Triton X-100, EDTA free Roche protease inhibitors, 1mM PMSF, DNase, 1mM BME. Process cells through microfluidizer for cell lysis. Centrifuge sample at 35000g at 4°C for 30 minutes. Load samples onto Talon metal affinity column for 60 min. Elute protein with 200 mM imidazole using a wash buffer consisting of 50 mM HEPES, pH 7.3, 300 mM NaCl, 5% glycerol, 0.03% Triton X-100, 1 mM BME. Change buffer composition to 20 mM HEPES, pH 7.3, 100 mM NaCl, 5% glycerol, 0.01% Triton X-100, 1 mM DTT with a G25 Sephadex desalting column. Bind protein by running solution through MonoS column with a buffer of 20 mM HEPES, pH 7.3, 5% glycerol, 0.01% Triton X-100, 1 mM DTT. Elute fractions with a salt gradient of 1 M NaCl. Separate protein according to size via a Superdex 200 column in a gel filtration buffer made of 20 mM HEPES, pH 7.3, 300 mM NaCl, 10% glycerol, 1 mM TCEP.

##### PIP4K2B

Codon optimized gene sequence encoding human PIP4K2B isoform 2 (Uniprot # P78356) was cloned into a pETM derived bacterial expression vector to create the following fusion protein: his6-SUMO3-GGGGG-PIP4K2B (1-416aa). Throughout the manuscript PIP4K2B is referred to as PIP4K. Recombinant PIP4K2B was expressed in BL21(DE3) Star *E. coli* (i.e. lack endonuclease for increased mRNA stability). Using 2-4 liters of Terrific Broth, bacterial cultures were grown at 37°C until OD_600_=0.6. Cultures were then shifted to 18°C for 1 hour to cool down. Protein expression was induced with 50 μM IPTG and bacteria expressed protein for 20 hours at 18°C before being harvested by centrifugation. For purification, cells were lysed into buffer containing 50 mM Na_2_HPO_4_ [pH 8.0], 400 mM NaCl, 0.4 mM BME, 1 mM PMSF (add twice, 15 minutes intervals), DNase, 1 mg/mL lysozyme using a microtip sonicator. Lysate was centrifuged at 16,000 rpm (35,172 x *g*) for 60 minutes in a Beckman JA-17 rotor chilled to 4°C. Lysate was circulated over 5 mL HiTrap Chelating column (GE Healthcare, Cat# 17-0409-01) that had been equilibrated with 100 mM CoCl_2_ for 1 hour, washed with MilliQ water, and followed by buffer containing 50 mM Na_2_HPO_4_ [pH 8.0], 400 mM NaCl, 0.4 mM BME. Recombinant PIP4K2B was eluted with a linear gradient of imidazole (0-500 mM, 8 CV, 40 mL total, 2 mL/min flow rate). Peak fractions were pooled, combined with 50 μg/mL of his6-SenP_2_ (SUMO protease), and dialyzed against 4 liters of buffer containing 25 mM Na_2_HPO_4_ [pH 8.0], 400 mM NaCl, and 0.4 mM BME for 16-18 hours at 4°C. Following overnight cleavage of the SUMO3 tag, dialysate containing his6-SUMO3, his6-SenP_2_, and GGGGG-PIP4K2B was recirculated for at least 1 hr over a 5 mL HiTrap(Co^+2^) chelating column. Flow-through containing GGGGG-PIP4K2B was then concentrated in a 30 kDa MWCO Vivaspin 6 before loading onto a Superdex 200 size exclusion column equilibrated in 20 mM HEPES [pH 7], 200 mM NaCl, 10% glycerol, 1 mM TCEP. In some cases, cation exchange chromatography was used to increase the purity of GGGGG-PIP4K2B before loading on the Superdex 200. In those cases, we equilibrated a MonoS column 20 mM HEPES [pH 7], 100 mM NaCl, 1 mM TCEP buffer. PIP4K2B (pI = 6.9) bound to the MonoS was resolved over a 10-100% linear gradient (0.1-1 M NaCl, 30 CV, 30 mL total, 1.5 mL/min flow rate). Peak fractions collected from the Superdex 200were concentrated in a 30 kDa MWCO Vivaspin 6 centrifuge tube and snap frozen at a final concentration of 20-80 μM using liquid nitrogen.

##### PLCδ-PH domain

The coding sequence of human PLCδ-PH domain (11-140aa) was expressed in BL21 (DE3) Star *E. coli* as a his_6_-SUMO3-(Gly)_5_-PLCδ (11-140aa) fusion protein. Bacteria were grown at 37°C in Terrific Broth to an OD_600_ of 0.8. Cultures were shifted to 18°C for 1 hour, induced with 0.1 mM IPTG, and allowed to express protein for 20 hours at 18°C before being harvested. Cells were lysed into 50 mM Na_2_HPO_4_ [pH 8.0], 300 mM NaCl, 0.4 mM BME, 1 mM PMSF, 100 μg/mL DNase using a microfluidizer. Lysate was then centrifuged at 16,000 rpm (35,172 x *g*) for 60 minutes in a Beckman JA-17 rotor chilled to 4°C. Lysate was circulated over 5 mL HiTrap Chelating column (GE Healthcare, Cat# 17-0409-01) charged with 100 mM CoCl_2_ for 1 hour. Bound protein was then eluted with a linear gradient of imidazole (0-500 mM, 8 CV, 40 mL total, 2 mL/min flow rate). Peak fractions were pooled, combined with SUMO protease (50 μg/mL final concentration), and dialyzed against 4 liters of buffer containing 50 mM Na_2_HPO_4_ [pH 8.0], 300 mM NaCl, and 0.4 mM BME for 16-18 hours at 4°C. Dialysate containing SUMO cleaved protein was recirculated for 1 hr over a 5 mL HiTrap Chelating column. Flow-through containing (Gly)_5_-PLCδ (11-140aa) was then concentrated in a 5 kDa MWCO Vivaspin 20 before being loaded on a Superdex 75 size exclusion column equilibrated in 20 mM Tris [pH 8.0], 200 mM NaCl, 10% glycerol, 1 mM TCEP. Peak fractions containing (Gly)_5_-PLCδ (11-140aa) were pooled and concentrated to a maximum concentration of 75 μM (1.2 mg/mL) before snap freezing with liquid nitrogen and storage at −80°C.

##### OCRL and DrrA

The coding sequence of human 5’-phosphatase OCRL (234-539aa of 901aa isoform) was expressed in BL21 (DE3) *E. coli* as a his6-MBP-(Asn)_10_-TEV-(Gly)_5_-OCRL fusion protein. DrrA/SidM (544-647aa of 647aa gene) derived from *L. pneomophila* was expressed in BL21 (DE3) *E. coli* as a his6-MBP-(Asn)_10_-TEV-(Gly)_5_-DrrA(544-647aa) fusion protein. For the proteins described above, bacteria were grown at 37°C in Terrific Broth to an OD_600_ of 0.8. Cultures were shifted to 18°C for 1 hour, induced with 0.1 mM IPTG, and allowed to express protein for 20 hours at 18°C before being harvested. Cells were lysed into 50 mM Na_2_HPO_4_ [pH 8.0], 300 mM NaCl, 0.4 mM BME, 1 mM PMSF, 100 μg/mL DNase using a microfluidizer. Lysate was then centrifuged at 16,000 rpm (35,172 x *g*) for 60 minutes in a Beckman JA-17 rotor chilled to 4°C. Lysate was circulated over 5 mL HiTrap Chelating column (GE Healthcare, Cat# 17-0409-01) charged with 100 mM CoCl_2_ for 1 hour. Bound protein was then eluted with a linear gradient of imidazole (0-500 mM, 8 CV, 40 mL total, 2 mL/min flow rate). Peak fractions were pooled, combined with TEV protease (75 μg/mL final concentration), and dialyzed against 4 liters of buffer containing 50 mM Na_2_HPO_4_ [pH 8.0], 300 mM NaCl, and 0.4 mM BME for 16-18 hours at 4°C. Dialysate containing TEV protease cleaved protein was recirculated for 1 hr over a 5 mL HiTrap Chelating column. Flow-through containing (Gly)_5_-protein was then concentrated in a 5 kDa MWCO Vivaspin 20 before being loaded on a Superdex 75 (10/300 GL) size exclusion column equilibrated in 20 mM Tris [pH 8.0], 200 mM NaCl, 10% glycerol, 1 mM TCEP. Peak fractions were pooled and concentrated before snap freezing in liquid nitrogen.

##### DrrA-OCRL

Chimeric 5’-phosphatase his6-MBP-(Asn)_10_-TEV-(Gly)_5_-DrrA(544-647aa)-(Gly)_5_-OCRL was expressed in BL21 (DE3) *E. coli*. For the proteins described above, bacteria were grown at 37°C in Terrific Broth to an OD_600_ of 0.8. Cultures were shifted to 18°C for 1 hour, induced with 0.1 mM IPTG, and allowed to express protein for 20 hours at 18°C before being harvested. Cells were lysed into 50 mM Na_2_HPO_4_ [pH 8.0], 300 mM NaCl, 0.4 mM BME, 1 mM PMSF, 100 μg/mL DNase using a microfluidizer. Lysate was then centrifuged at 16,000 rpm (35,172 x *g*) for 60 minutes in a Beckman JA-17 rotor chilled to 4°C. Lysate was circulated over 5 mL HiTrap Chelating column charged with 100 mM CoCl_2_ for 1 hour. Bound protein was then eluted with a linear gradient of imidazole (0-500 mM, 8 CV, 40 mL total, 2 mL/min flow rate). Peak fractions were pooled, combined with TEV protease (75 μg/mL final concentration), and dialyzed against 4 liters of buffer containing 50 mM Na_2_HPO_4_ [pH 8.0], 300 mM NaCl, and 0.4 mM BME for 16-18 hours at 4°C. Dialysate containing TEV protease cleaved protein was recirculated for 1 hr over a 5 mL HiTrap Chelating column. Flow-through containing (Gly)_5_-DrrA(544-647aa)-(Gly)_5_-INPP5E or (Gly)_5_-DrrA(544-647aa)-(Gly)_5_-OCRL were then buffer exchanged into 20 mM HEPES pH 7, 100 mM NaCl, 1 mM DTT using a HiPrep 26/10 desalting column (GE Healthcare, Cat# 17-5087-01). DrrA-OCRL or DrrA-INPP5E were then loaded onto a 1 mL MonoS (5/50 GL) cation exchange column (GE Healthcare, Cat# 17-5168-01) equilibrated in 20 mM HEPES pH 7, 100 mM NaCl, 1 mM DTT. DrrA-OCRL and DrrA-INPP5E were separated from impurities by applying a linear salt gradient (0.1-1 M NaCl) over 45 CV (45 mL total). Both DrrA-OCRL and DrrA-INPP5E eluted from the MonoS column in the presence of 250-300 mM NaCl. Peak fractions were pooled and concentrated in a 10 kDa MWCO Vivaspin 6 before being loaded on a Superdex 75 (10/300 GL) size exclusion column equilibrated in 20 mM Tris [pH 8.0], 200 mM NaCl, 10% glycerol, 1 mM TCEP. Peak fractions were pooled and concentrated before snap freezing in liquid nitrogen.

##### Sortase mediated peptide ligation

All lipid sensors and catalytic domains were labeled on a N-terminal (Gly)_5_ motif using sortase mediated peptide ligation (Ton-That et al. 1999; Guimaraes et al. 2013). We devised a novel approach for chemically modifying an LPETGG peptide with fluorescent dyes, which we then conjugated to our protein of interest. The LPETGG peptide was synthesized to >95% purity by ELIM Biopharmaceutical (Hayward, CA) and labeled on the N-terminal amine with N-Hydroxysuccinimide (NHS) fluorescent dye derivatives (e.g. NHS-Alexa488). This was achieved by combining 10 mM LPETGG peptide, 15 mM NHS-Alexa488 (or other fluorescent derivatives), and 30 mM Triethylamine (Sigma, Cat# 471283) in anhydrous DMSO (Sigma, Cat# 276855). This reaction was incubated overnight in the dark at 23°C before being stored in a −20°C freezer. Prior to labeling (Gly)_5_ containing proteins, unreacted NHS-Alexa488 remaining in the LPETGG labeling reaction was quenched with 50 mM *Tris*(hydroxymethyl)aminomethane (Tris) [pH 8.0] buffer for at least 6 hours. Complete quenching of unreacted NHS-Alexa488 was verified by the inability to label (Gly)_5_ containing proteins in the absence of a Sortase.

When labeling (Gly)_5_ containing proteins with the fluorescently labeled LPETGG peptide we typically combined the following reagents: 50 mM Tris [pH 8.0], 150 mM NaCl, 50 μM (Gly)_5_-protein, 500 μM Alexa488-LPETGG, and 10-15 μM His_6_-Sortase (Δ57; lacks first 57 amino acids). This reaction mixture was incubated at 16-18°C for 16-20 hours, before buffer exchange with a G25 Sephadex column (e.g. PD10 or NAP5) to remove majority of dye and dye-peptide. The his_6_-Sortase was then captured on NiNTA agarose resin (Qiagen) and unbound, labeled protein was separated from remaining fluorescent dye and peptide using a Superdex 75 or Superdex 200 size exclusion column (24 mL bed volume).

##### Preparation of small unilamellar vesicles

The following lipids were used to generated small unilamellar vesicles (SUVs): 1,2-dioleoyl-sn-glycero-3-phosphocholine (18:1 DOPC, Avanti # 850375C), L-α-phosphatidylinositol-4-phosphate (Brain PI(4)P, Avanti Cat# 840045X), L-α-phosphatidylinositol-4,5-bisphosphate (Brain PI(4,5)P_2_, Avanti # 840046X), 1,2-dipalmitoyl-sn-glycero-3-phosphoethanolamine-N-(lissamine rhodamine B sulfonyl) (16:0 Liss Rhod PE, Avanti # 810158C), 1,2-dioleoyl-*sn-*glycero-3-phospho-L-serine (18:1 DOPS, Avanti # 840035C). In the main text, 16:0 Liss Rhod PE is referred to as Rhod PE. Lipids were purchased as single use ampules containing between 0.1-5 mg of lipids dissolved in chloroform. Brain PI(4)P and PI(4,5)P_2_ were purchased as 0.25 mg/mL stocks dissolved in chloroform:methanol:water (20:9:1). To make liposomes, 2 μmoles total lipids are combined in a 35 mL glass round bottom flask containing 2 mL of chloroform. Lipids are dried to a thin film using rotary evaporation with the glass roundbottom flask submerged in a 42°C water bath. After evaporating all the chloroform, the round bottom flask was flushed with nitrogen gas for at least 30 minutes. Resuspend lipid film in 2 mL of PBS [pH 7.2], making a final concentration of 1 mM total lipids. All lipid mixtures expressed as percentages (e.g. 98% DOPC, 2% PI(4)P) are equivalent to molar fractions. For example, a 1 mM lipid mixture containing 98% DOPC and 2% PI(4)P is equivalent to 0.98 mM DOPC and 0.02 mM PI(4)P. To generate 30-50 nm SUVs, 1 mM total lipid mixtures were extruded through a 0.03 μm pore size 19 mm polycarbonate membrane (Avanti #610002) with filter supports (Avanti #610014) on both sides of the PC membrane. Hydrated lipids at a concentration of 1 mM were extruded through the PC membrane 11 times.

##### Preparation of supported lipid bilayers

Supported lipid bilayers are formed on 25×75 mm coverglass (IBIDI, #10812). Coverglass is first cleaned with 2% Hellmanex III (Fisher, Cat#14-385-864) heated to 60-70°C in a glass coplin jar. Incubate for at least 30 minutes. Wash coverglass extensively with MilliQ water and then etched with Pirahna solution (1:3, hydrogen peroxide:sulfuric acid) for 10-15 minutes the same day SLBs were formed. Etched coverglass, in water, is rapidly dried with nitrogen gas before adhering to a 6-well sticky-side chamber (IBIDI, Cat# 80608). Form SLBs by flowing 30 nm SUVs diluted in PBS [pH 7.2] to a total lipid concentration of 0.25 mM. After 30 minutes, IBIDI chambers are washed with 5 mL of PBS [pH 7.2] to remove non-absorbed SUVs. Membrane defects are blocked for 15 minutes with a 1 mg/mL beta casein (Thermo FisherSci, Cat# 37528) diluted in 1x PBS [pH 7.4]. Before use as a blocking protein, frozen 10 mg/mL beta casein stocks were thawed, centrifuged for 30 minutes at 21370 x *g*, and 0.22 μm syringe filtered. After blocking SLBs with beta casein, membranes were washed again with 1mL of PBS, followed by 1 mL of kinase buffer before TIRF-M.

##### Kinetics measurements of PIP lipid phosphorylation

The kinetics of PI(4)P phosphorylation was measured on SLBs formed in IBIDI chambers and visualized using TIRF microscopy. Reaction buffer contained 20 mM HEPES [pH 7.0], 150 mM NaCl, 1 mM ATP, 5 mM MgCl_2_, 0.5 mM EGTA, 20 mM glucose, 200 μg/mL beta casein (ThermoScientific, Cat# 37528), 20 mM BME, 320 μg/mL glucose oxidase (Serva, #22780.01 *Aspergillus niger*), 50 μg/mL catalase (Sigma, #C40-100MG Bovine Liver), and 2 mM Trolox (UV treated, see methods below). Perishable reagents (i.e. glucose oxidase, catalase, an Trolox) were added 5-10 minutes before image acquisition. For all experiments, we monitored the change in PI(4)P or PI(4,5)P_2_ membrane density using a solution concentrations of 20 nM Alexa647-DrrA(544-647) or 20 nM Alexa488-PLCδ, respectively. We calculated a density of PIP lipids (lipids/μm^2^) assuming a footprint of 0.72 nm^2^ for DOPC lipids (Galush, Nye, and Groves 2008; Vacklin, Tiberg, and Thomas 2005).

##### Single molecule cell lysate assay

Genes encoding hPIP5KA, hPIP5KB, hPIP5KG, zPIP5KA, yMss4, and hPIP4K2B were cloned into lentiviral expression vectors containing a SFFV promoter to drive expression of N-terminal mNeonGreen fusion proteins in mammalian cells. HEK293T cells were transfected with 15 μg of plasmid DNA encoding mNG tagged lipid kinases, plus 30 μg polyethylenimine (PEI) diluted into 0.5 mL Opti-MEM. Prior to transfection, HEK293T cells were grown to a confluency of 50-60% in 10 cm dishes in DMEM GlutMax media (ThermoFisher, Cat #10566016) containing 10% FBS and penicillin/streptomycin. After 20-24 hours for transfection, adherent HEK293T grown in 10 cm dishes were washed with 5 mL 1x PBS [pH 7.4]. After vacuum aspiration of the PBS, cells were incubated in 1 mL of CellStripper (Corning, Cat# 25-056-Cl) for 10 minutes at room temperature. Detached cells were resuspended in 9 mL of PBS and transferred to a 15 mL conical tube. Cells were centrifuged for 5 minutes at 500 rcf at 4°C. The cell pellet was resuspended in 1 mL of PBS, transferred to a microcentrifuge tube, and centrifuged for 3 minutes at 500 rcf. After removing PBS by vacuum aspiration, cell pellets were resuspended in 0.6 mL of lysis buffer containing 20 mM HEPES pH 7, 150 mM NaCl, 5 mM MgCl_2_, Sigma protease inhibitor (Cat# P3840, 1% vol./vol. final), Sigma PPtase inhibitor 2 (0.5% vol./vol. final), Sigma PPtase inhibitor 3 (0.5% vol./vol. final), 50 mM NaF, 15 μg/mL benzamidine, and 1mM PMSF. Microtip sonication was used to rupture transfected cell on ice using the following program: 20% amplitude, 2 sec ON and 20 sec OFF for 15 cycles. Cell lysate was centrifuged for 45 min at 21300 rcf at 4°C. Following centrifugation, 75% of the supernatant was transferred to new 1.7 mL microcentrifuge tube. The clarified lysate was then mixed 4:1 with lysis buffer containing 50% glycerol (vol./vol.). This resulted in lysate containing a final glycerol concentration of 10% (vol./vol.). Cell lysate was aliquoted and flash frozen in liquid nitrogen. We did not observe any difference in the quality of the mNG labeled protein comparing fresh versus freeze thawed cell lysate.

To determine the concentration of mNG tagged lipid kinase in HEK293T cell lysate we generated a 2-fold serial dilution of bacterially purified mNeonGreen diluted in 1x PBS [pH 7.4] and 0.1% NP-40 detergent. The fluorescence intensity was measured using a BioTek 96-well format using a plate reader to generate a standard curve for fluorescence intensity as a function of mNG concentration (**Figure S1A**). mNG was excited with a 500 nm light using a 500/10nm bandpass filter. The emission was monitored at 517 nm with high photomultiplier tube (PMT) sensitivity.

##### Quantitative fluorescence microscopy using supported lipid bilayer standards

We measure the membrane surface density of Alexa647-PIP5KB (**Figure 2**) using previously described methods (Galush, Nye, and Groves 2008). In brief, we titrated the molar fraction of Atto655-1,2-dipalmitoyl-sn-glycero-3-phosphoethanolamine (Atto655-DPPE; Atto-TEC, Cat# AD 655-151) against DOPC lipids. Small unilamellar vesicles (SUVs) were formed using microtip sonication. We used TIRF microscopy to measure the fluorescence intensity of supported lipid bilayers containing varying concentrations of Atto655-DPPE. These values we used to generated a standard curve that was used to calculate the surface density of membrane bound Alexa647-PIP5KB. In order to compare the fluorescence intensity of Alexa647-PIP5KB to Atto655-DPPE, we calculated the scaling factor which accounts for the difference in molecular brightness of the two fluorophores measured on the same microscope with identical camera settings, laser power, filters, and optics. To determine the scaling factor we measured the fluorescence intensity of solutions containing identical concentrations of either Alexa647-PIP5KB or SUVs containing the same molar concentration of Atto655-DPPE (**Figure S1**). These solution were added to an imaging chamber passivation with a supported membrane (95% DOPC and 5% DOPS) in order to prevent non-specific absorption of Alexa647-PIP5KB and SUVs containing Atto655-DPPE. To measure the solution intensity of the Alexa647-PIP5KB and Atto655-DPPE containing samples, we choose a z-axis imaging plane that was 5 μm above the glass surface and acquired at least 20 fluorescence measurement for each solution concentration. These fluorescence intensity values were averaged and plotted as a function of the fluorophore solution concentration. The scaling factor was calculated by dividing the slope of Alexa647-PIP5KB plot by the slope of the Atto655-DPPE plot.

##### Surface density calibration of mNG-PIP5KB

Densities of mNG-PIP5KB were estimated using a surface density calibration curve of mNG attached to the lipid bilayer. Supported lipid bilayer containing 96% DOPC and 4% Ni-NTA-DOGS by molar percent was incubated with His6-mNG with concentrations ranging from 0.1 nM to 10 nM in 20 mM HEPES [pH 7.0], 150 mM NaCl, 200 μg/mL beta casein, 5 mM BME for 30 minutes. The chambers were rinsed with 1 mL of buffer and a calibration curve was established between the TIRF average intensity and surface densities measured by fluorescence correlation spectroscopy (FCS) of mNG on the membrane using previously described methods (Chung et al. 2019; 2018). The same calibration curve was used to estimate the surface density of mNG-PIP5KB. The kinetic measurement were performed on supported lipid bilayer containing 96% DOPC and 4% PI(4)P by molar percent, with the presence of 20 nM Alexa488-PLCδ to monitor the change of PI(4,5)P_2_. The total PIP lipid was calculated to be at 55555 lipids/μm^2^. The start and end of the reaction were approximated to have 0 and 55555 lipids/μm^2^ of PI(4,5)P_2_, respectively. Images were acquired at a 2 sec interval. PI(4,5)P_2_ and dPI(4 5)P_2_ mNG-PIP5KB surface density was calculated for each time point. The 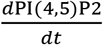 for each time point was obtained by using the slope of linear regression of PI(4,5)P_2_ level change in a +2 sec to −2 sec time range.

##### Feedback analysis of PIP5K

Alexa488-PLCδ intensity was measured from TIRF images acquired at a 2 sec interval. The reaction coordinate (x) for each time point was calculated by normalizing the start intensity to 0 and end intensity to 1. The 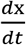 for each time point was obtained by using the slope of linear regression of reaction coordinate change in a +2 sec to −2 sec time range.

##### Chromium patterned glass coverslips

25 x 75 mm No. 1.5 thickness glass coverslips were cleaned in acetone by sonication, then washed with MilliQ water extensively. The coverslips were dried by nitrogen gas then baked on 120°C hot plate for 5 minutes. S1805 positive photoresist were spin coated on the coverslips by spinning for 2 seconds at 500 rpm (ACL 440) then for 30 seconds at 4111 rpm (ACL 3900). The photoresist on the edge of the coverslips were removed by cotton swap soaked with acetone, then baked on 120°C hot plate for 1 minute. Mask with desired pattern was mounted on an OAI Series 200 Aligner. The photoresist coated coverslip was exposed for 0.6 sec with UV power around 30 mJ, then developed with MicroPosit MF-321 Liquid Developer for 40 sec with mild shaking. The developed coverslips were rinsed with water and dried with nitrogen gas. ~9 nM thick chromium was subsequently deposited on the coverslips using an electron beam evaporator at 1×10^-6^ torr. Finally, photoresist is lifted from chromium patterned glass substrates by bath sonication in Dow Electronic Materials MicroPosit Remover 1165 for 10 minutes for 2 times, then washed with water.

##### Microscope hardware and imaging acquisition

Single-molecule imaging experiments were performed on an inverted Nikon Eclipse Ti and Ti2 microscopes using a 100x Nikon objective (1.49 NA) oil immersion TIRF objective. The macroscopic spatial patterning of PIP compositional patterns and the chromium patterned SLB were visualized using a 60x Apo TIRF oil immersion objective (1.45 NA). We manually control the x-axis and y-axis positions using an ASI stage and joystick. All images were acquired using either a iXon Ultra or iXion Life 897 EMCCD camera (Andor Technology Ltd., UK). Fluorescently labeled proteins were excited with either a 488 nm, 561 nm, or 637 nm diode laser (OBIS laser diode, Coherent Inc. Santa Clara, CA) controlled with either a Solemere (Nikon Ti) or Vortran (Nikon Ti2) laser drive with acousto-optic tunable filters (AOTF) control. The power output measured through the objective for single particle imaging was 1-3 mW. For dual color imaging of spatial PIP lipid patterns on SLBs, samples were excited with 0.2-0.5 mW 488 nm and 0.2-0.5 mW 637 nm light, as measured through the objective. Excitation light was passed through the following dichroic filter cubes before illuminating the sample: (1) ZT488/647rpc and (2) ZT561rdc (ET575LP) (Semrock). Fluorescence emission was detected on an ANDOR EMCCD camera position after a Sutter emission filter wheel housing the following emission filters: ET525/50M, ET600/50M, ET700/75M (Semrock). All experiments were performed at room temperature (23°C). Microscope hardware was controlled using both Micro-Manager v4.0 (Edelstein et al. 2010) and Nikon NIS elements.

##### Single particle tracking

Fluorescent particle detection and tracking was performed using the ImageJ/Fiji TrackMate plugin (Jaqaman et al. 2008). Image stacks containing ~1000 16-bit image stacks in the form of a .nd2 file were loaded in ImageJ. Image sequences were cropped to 400×400 pixels in order to minimize differences in field illumination caused by TIRF illumination. Using the LoG detector option, particles were identify based on brightness and their signal-to-noise ratio. After identifying the position of all fluorescent particle, we used the LAP tracker to generate particle trajectories that followed molecular displacement as a function of time. Particle trajectories were then filtered based on Track Start (removed trajectories that began in first frame), Track End (removed trajectories present in last frame), Duration (removed trajectories ≤ 2 frames), Track displacement (removed immobilized particles), and X - Y location (removed particles near the edge of the images). Trajectories were filtered to remove 1-5% of particles that were immobilized throughout the image sequence. The TrackMate output files were analyzed using custom MATLAB scripts to calculate the single molecule dwell times and diffusion coefficients.

Step size distribution of single particle trajectories were plotted in MATLAB and Prism as frequency versus step size (μm). For all analysis presented in this manuscript, the bin size for the step size distribution equals 0.01 μm. For curving fitting, the step-size distribution were plotted as probability density versus step-size (μm). This was achieved by dividing the frequency distribution by the bin size (0.01 μm). Probability density versus step size plots was fit to the following one- or two-species distributions:

Single species model:

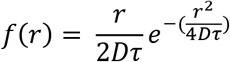
Two species model:

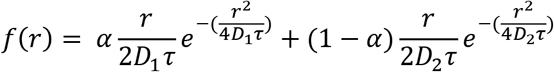

Variables are defined as the D1=diffusion coefficient species 1 (μm^2^/sec), D2=diffusion coefficient species 2 (μm^2^/sec), alpha (τ_1_ = % of species 1, r = step size (μm), τ_2_ = time interval between steps (sec). Final step size distribution plots were generated in PRISM graphing software and using the following equations: (1 species model): f(x) = x/(2*D1*t)*exp(-(x^2/(4*D1*t))), (2 species model): f(x) = alpha*(x/(2*D1*t)*exp(-(x^2/(4*D1*t))))+(1-alpha)*(x/(2*D2*t)*exp(-(x^2/(4*D2*t)))).

To calculate the dwell times for membrane bound lipid kinases we sorted into a cumulative distribution frequency (CDF) plot with the frame interval as the bin (e.g. 50 ms). The log_10_(1-CDF) is then plotted against the dwell time and fit to a single or double exponential.

Single exponential model:

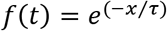

Two exponential model:

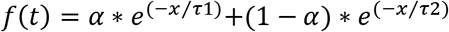

Fitting procedure initiated with a single exponential. In cases of a low quality single exponential fit, a maximum of two species model was used. For double exponential fit, alpha represents the fraction of fast dissociating molecules characterized by τ_1_.

##### Image analysis, curve fitting, and statistics

Image analysis was performed on ImageJ and MatLab. Curve fitting was performed using MatLab and Prism 8 (GraphPad). Single molecule dwell time, step size, and molecular brightness distributions presented in this manuscript represent combined data from 3 technical replicates with 2-3 movies acquired from multiple fields of view for each experimental condition. Dwell time distributions and curve fits were generated with n = 1000-3000 particle trajectories (**Figures 2B, 3C-3E**). Step size distribution plots and curve fits represent 10,000-30,000 measured displacements (**Figures 1F, 2C, 3F** and **Figure 3–figure supplement 1**). Single molecule brightness distribution plots were generated from analyzing n = 15000-75000 fluorescent particles (**Figures 1E** and **Figure 1–figure supplement 2**).

